# Age-dependent sexual dimorphism in the adult human gut microbiota

**DOI:** 10.1101/646620

**Authors:** Xiuying Zhang, Huanzi Zhong, Yufeng Li, Zhun Shi, Zhe Zhang, Xianghai Zhou, Huahui Ren, Shanmei Tang, Xueyao Han, Yuxiang Lin, Dan Wang, Fangming Yang, Chao Fang, Zuodi Fu, Lianying Wang, Shida Zhu, Yong Hou, Xun Xu, Huanming Yang, Jian Wang, Karsten Kristiansen, Junhua Li, Linong Ji

## Abstract

A decade of studies has established the importance of the gut microbiome in human health. In spite of sex differences in the physiology, lifespan, and prevalence of many age-associated diseases, sex and age disparities in the gut microbiota have been little studied. Here we show age-related sex differences in the adult gut microbial composition and functionality in two community-based cohorts from Northern China and the Netherlands. Consistently, women harbour a more diverse and stable microbial community across broad age ranges, whereas men exhibit a more variable gut microbiota strongly correlated with age. Reflecting the sex-biased age-gut microbiota interaction patterns, sex differences observed in younger adults are considerably reduced in the elderly population. Our findings highlight the age- and sex-biased differences in the adult gut microbiota across two ethnic population and emphasize the need for considering age and sex in studies of the human gut microbiota.

## Introduction

Along with rapid socio-economic and lifestyle changes during the past decades in China, the prevalence of chronic non-communicable diseases (NCDs) has increased dramatically ^1,2^. Accumulating evidence from epidemiology studies has revealed age-related sex differences in life expectancy, and in risk, course, and outcomes of NCDs ^3,4^. Paradoxically, women live longer, but are predisposed to higher incidences and worse outcomes of certain diseases than age-matched men in late life, such as cardiometabolic disorders and Alzheimer’s disease ^5–8^.

In spite of increasing evidence pointing to associations between the gut microbiota and NCDs ^9–12^, much remains unknown in relation to sexual dimorphism in the gut microbiota as well as possible interactions with sex hormones, ageing, and sex- and age-specific health conditions. Two pioneer mice studies have demonstrated that a sexual dimorphic gut microbiota may bidirectionally interact with host testosterone levels and further influence the lifetime risk of autoimmune diseases ^13,14^. Using 89 inbred mice strains, Elin *et al.* have further elaborated on genetics-dependent sex differences in the gut microbial composition ^15^. However, information on sexual dimorphism in the human gut microbiota is very limited, but a recent study reported on sex-specific differences, especially in gut resistome profiles, in a large-scale Dutch cohort (LifeLine DEEP cohort, LLD) ^16^. In addition, the interplay between the gut microbiota and sex hormones, and sex- and age-specific health conditions so far has only been investigated in women with small sample sizes ^17,18^.

In this study, we systematically investigated gut microbial characteristics and their associations with sex hormones and additional extensive host metadata on 2,338 adults (26-76 years) from a Han Chinese population-based cohort established in the Pinggu (PG) district, Beijing, the PG cohort. Taking advantages of the same ethnic background and shared geography and by removing samples from adults taking commonly used medicines for metabolic disorders, we have largely avoided the reported metagenomic confounding effects related to differences in host genetics, lifestyle, dietary patterns, and medicine use in this cohort ^19–21^. Thereby, we uncovered striking age-dependent sex differences in the gut microbial composition and functionality and replicated our main findings in the LLD cohort. Compared to men, women overall harboured a more diverse microbiota, showing weak associations with age, menopause, and menopause related declines in sex hormones and health conditions, exhibiting minor changes across different age groups. By contrast, the gut microbiota of men tended to be more plastic with marked differences between different age groups, displaying strong associations with age, alcohol intake, smoking, and testosterone. Due to age- and sex-biased changes in the gut microbiota, the magnitude of sex-associated gut microbial differences decreased in the elderly (above 50 years) compared to the younger (below 50 years) adults.

Our findings provide novel insights into the characteristics of the sex-biased adult gut microbiota and correlations with age, sex hormones, lifestyle, and host health conditions. Further longitudinal studies are warranted to investigate the underlying mechanisms governing the different sex-dependent developmental trajectories of gut microbial communities and the potential impact on lifespan and health.

## Results

### The PG cohort

The PG cohort is a Chinese Han population-based prospective cohort established in the Pinggu district of Beijing in North China to study how environmental factors, diet, host physiology and behaviour, and the gut microbiota might contribute to or be associated with the growing NCD epidemic. From this cohort, metagenomic data of faecal samples from 2,338 individuals aged 26-76 years were analysed with time-matched clinical measures for phenotyping, sex hormone levels, and host metabolic status, supplemented with questionnaire data covering lifestyle, diet, intake of drugs, and female gynaecological information **(Methods, Fig. 1a, Supplementary Table 1**).

**Fig 1.**
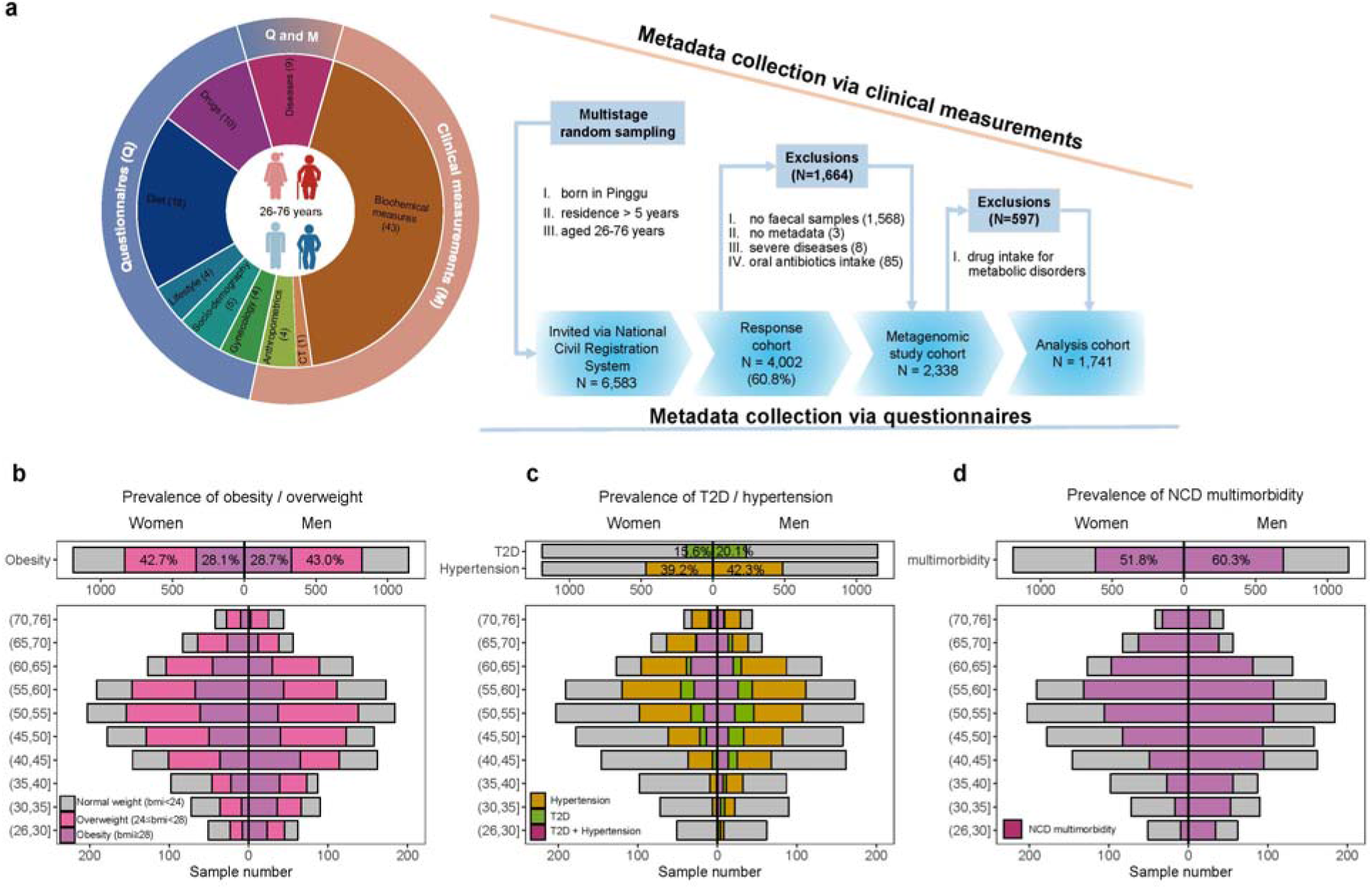
Overview of the Pinggu cohort. **a,** In the PG cohort, 4,002 of 6,538 invited individuals drawn by multistage stratified sampling responded. 2,338 individuals meeting the additional criteria were included in metagenomic study, of which 1,741 free of drugs for treating diabetes, hypertension or dyslipidemia constituted the final analysis cohort. The study collected 98 phenotypic factors in total through clinical measurements (M): biochemical measures in blood / urine samples (n=43), anthropometric measures (n=4), computed tomography (CT) examination (n=1); through questionnaires (Q): female gynecological information (n=4), socio-demography (n=5), lifestyle (n=4), diet (n=18) and drugs (n=10), and diseases (n=9) based on M and Q. The summary statistics of all factors is shown in **Supplementary Table S1**. **b-d,** Population pyramids showing the age-sex distribution of the PG metagenomic study cohort (N=2,338) and the prevalence of obesity / overweight **(b)**, of type 2 diabetes (T2D) and hypertension **(c),** and of non-communicable diseases (NCD) multimorbidity **(d)** for women (left) and men (right) within each age group. NCD multimorbidity is defined as the presence of two or more of the six chronic metabolic conditions, including obesity, dyslipidaemia, fatty liver disease, hyperuricemia, T2D and hypertension **(Methods)**.

The PG cohort exhibited a greater prevalence of obesity than a recently reported Shanghai cohort (**Fig. 1b**, 28.5% vs 12.4%) ^22^, as well as high prevalence of type 2 diabetes (**Fig. 1c**, 15.6% for women and 20.1% for men), and NCD multimorbidity (51.8% for women and 60.3% for men), defined as the presence of two or more of six chronic metabolic conditions, including obesity, dyslipidaemia, hyperuricemia, hypertension, T2D, and fatty liver disease (FLD) diagnosed by a liver-to-spleen (L/S) attenuation ratio ≤ 1.1 using computerized tomography (CT) scanning ^23^ (**Methods, Fig. 1d, Supplementary Table 1)**. 597 NCD patients (25.53% of the 2,338 individuals) reported upon collection of faecal samples the use of drugs known to improve blood pressure, glucose, and lipids (**Supplementary Table 2)**. In agreement with previous studies ^19,24,25^, drugs such as biguanides, glucosidase inhibitors (GIs) and statins, showed significant effects on the composition of the gut microbiota, especially when taken in combination (**Methods, PERMANOVA,** adjusted *P* < 0.05, **Supplementary Table 2**). To avoid the confounding effects of drugs on host biochemical levels, gut microbiota, and the possible mutual interactions, we excluded data from these 597 NCD patients and established an analysis cohort of 1,741 adults with no reported drug treatment for further analyses (**Supplementary Table 3**).

### Sex differences in gut microbial composition and functionality

To investigate the importance of covariates in the PG cohort, PERMANOVA was applied for host phenotypes collected through clinical measurements and questionnaires **(Fig. 1a, Methods).** In the entire analysis cohort, 42 factors were identified as significant covariates impacting on the gut microbiota, including sex, age, serum triglyceride (TG), uric acid (UA), testosterone levels, waist-to-hip ratio (WHR), body mass index (BMI), and male characteristic lifestyles, including intake of alcohol and smoking (**PERMANOVA,** adjusted *P* < 0.05, **Fig. 2a, Supplementary Table 4-6**). Interestingly, sex explained the largest gut microbial variance in the PG cohort **(Fig.2a)**, whereas sex was found to rank lower than other commonly identified covariates such as TG, UA and BMI in previous large-scale studies ^20,26,27^.

**Fig 2.**
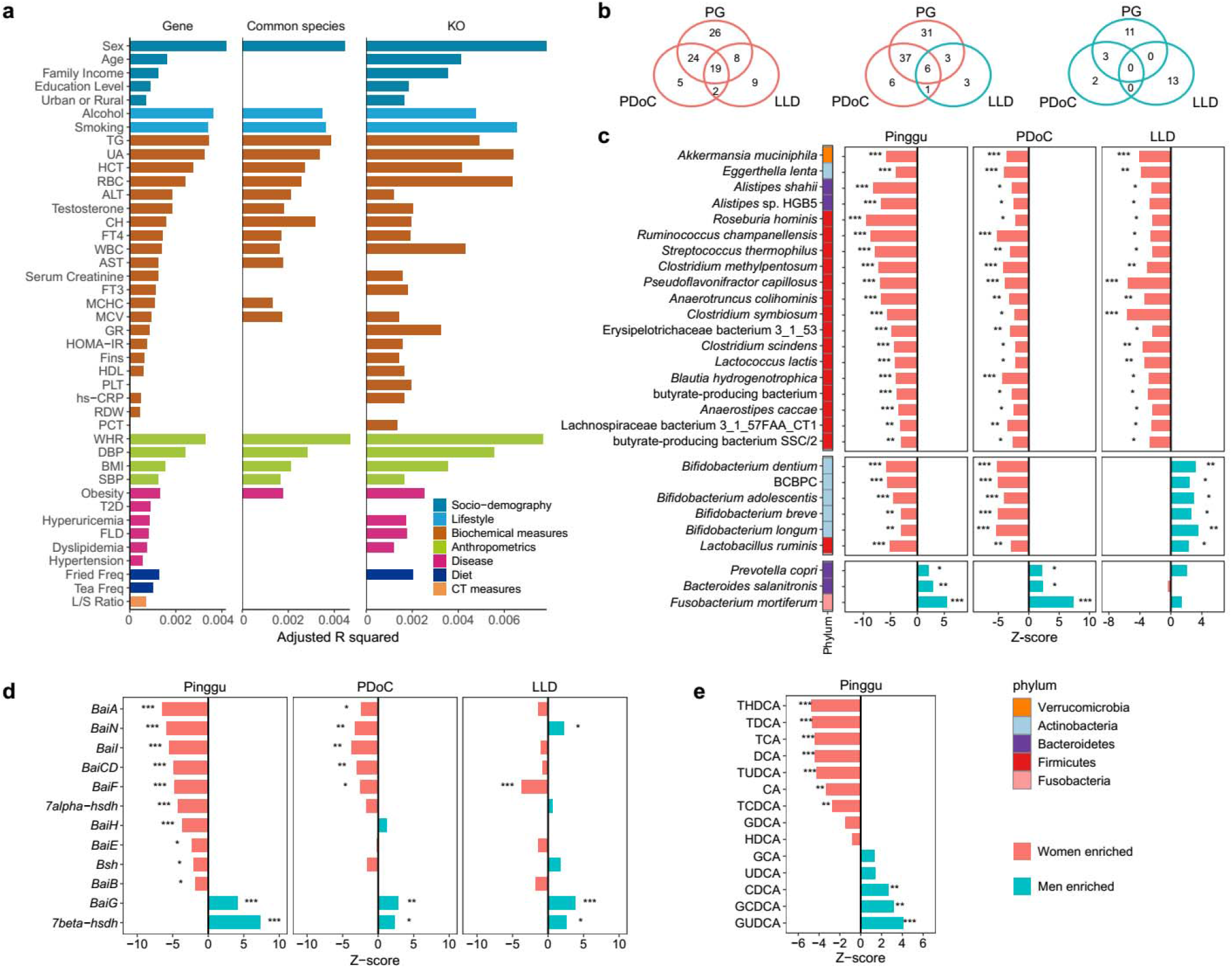
Sex differences in gut microbial composition and functionality. **a,** Horizontal bars showing the amount of inferred variance (adjusted R-squared) explained by each identified covariate as determined by PERMANOVA with Bray-Curtis (BC) dissimilarities at the gene (left), common species (middle) and KEGG Orthology (KO, right) level. Metadata categories are indicated by colors and ranked by the highest explained variation in the respective category. Only statistically significant covariates with adjusted *P* < 0.05 using Benjamini and Hochberg (BH) method are shown. **b,** Venn diagrams showing the overlap of sex-dependent differentially abundant species between the PG cohort (up), the published datasets of Chinese adults (PDoC, left) and the LifeLines DEEP cohort (LLD, right). Red circles for women enriched species, and blue circles for men enriched species. Wilcoxon rank test, BH adjusted *P* < 0.05. **c,** Selected sex-dependent differentially abundant common species in the PG cohort (left), the published datasets of Chinese adults (middle), and the LLD cohort (right). The colored bar indicates phylum-level taxonomy of each species. BPBPC, represents *Bifidobacterium catenulatum-Bifidobacterium pseudocatenulatum* complex. **d,** Enrichment of relative abundance of genes involved in bile acid transformation and transport between the two sexes in the three datasets. **e**, Enrichment of relative concentrations of serum bile acids in a subgroup including 424 age-, sex-, and BMI-matched PG individuals. Wilcoxon rank test (**c-e**), Z-scores shown as horizontal bars indicate the enrichment direction between sexes, red for women enriched and blue for men enriched. * BH adjusted *P* < 0.05; ** adjusted *P* < 0.01; *** adjusted *P* < 0.001.

To characterize sex-associated differences in the gut microbiota, we performed comparisons of the metagenomes between women and men in the PG cohort, and furthermore, in two independent published shotgun metagenomic datasets of Chinese ^9–12, 25^ (n=876) and Dutch adults ^26^ (n=1,135, the LLD cohort) for validation purposes **(Methods)**. We uncovered that women in the PG cohort showed greater microbial α diversity at the gene, species, and KEGG Orthology (KO) level than men (*P* < 0.05, **Extended Data Fig. 1a**). These findings were consistently replicated in the published Chinese datasets, and in the LLD datasets with relatively smaller sex differences compared to two Chinese datasets, despite substantial differences in microbial diversity, Bacteroidetes to Firmicutes (B / F) ratio and enterotypes between Chinese and Dutch adults (**Extended Data Fig. 1-2, Supplementary Table 7**) ^28,29^.

Out of 151 common species, 91 differed significantly in abundance between sexes, and over half (77) were enriched in women (adjusted *P* < 0.05, **Fig. 2b, Supplementary Table 7**). Specifically, 19 species were significantly enriched in both Chinese and Dutch adult women, including *Akkermansia muciniphila, Eggerthella lenta, Alistipes shahii* and 15 Firmicutes species including *Ruminococcus champanellensis, Clostridium scindens, C. methylpentosum* and *C. symbiosum* (adjusted *P* < 0.05, **Fig. 2b-c, Supplementary Table 7**). Several species from *Bifidobacterium*, a main probiotic genus, were significantly enriched in Chinese women, but were by contrast enriched in men in the LLD cohort (**Fig. 2b-c)**. In addition, a significant enrichment of *Fusobacterium mortiferum, Prevotella copri* and *Bacteroides salanitronis* in men was observed specifically in Chinese (**Fig. 2b-c**). Reflecting the enrichment of a variety of Firmicutes bacteria capable of transforming bile acid (BA) in adult PG women ^30,31^, most bile acid inducible genes were also significantly enriched in women, except for the genes encoding the 7-beta-hydroxysteroid dehydrogenases (7-β-HSDH) and the bile acid transporter (Bai G), which were enriched in adult men (adjusted *P* < 0.05, **Fig. 2d**). In line with a previous study in a healthy Chinese population ^32^, we observed significant sex differences in serum BA profiles in a subgroup of 424 age-, sex- and BMI-matched PG individuals. Thus, we observed higher relative levels of cholic acid (CA) and CA-derived secondary BAs (deoxycholic acid (DCA) and hyodeoxycholic acid (HDCA)) in women, contrasted by higher levels of chenodeoxycholic acid (CDCA) and ursodeoxycholic acid (UDCA), and a larger total BA pool in men **(Fig. 2e, Extended Data Fig. 3, Supplementary Table 8**).

In summary, our results revealed substantial population independent and dependent sex differences in the gut microbial composition and functional capacity.

### Sex-biased host phenotype-gut microbiota associations

Substantial sex differences were also observed in host physiology, dietary patterns, and lifestyle, with men showing particularly higher prevalence of smoking (67.61% vs. 1.01%) and alcohol intake (80.7% vs. 13.65%) than women **(Extended Data Fig. 4a-d, Supplementary Table 3)**. Moreover, as expected postmenopausal women exhibited much lower levels of sex hormone and worse metabolic conditions, whereas elderly men (above 50 years of age) compared with younger men unexpectedly exhibited higher testosterone level and lower levels of obesity-related clinical parameters (adjusted *P* < 0.05, **Extended Data Fig. 4e-h)**.

Considering the sex disparity in the gut microbiota, host physiology, and behavior, we next conducted PERMANOVA analyses in each sex, investigating whether different host phenotype-gut microbiota association patterns characterised women and men. Surprisingly, many sex-biased covariates, including several routine blood parameters, education level, family income, and frequency in consumption of fried food and tea, identified in the analysis of the entire cohort (adjusted *P* < 0.05, **Fig. 2a**), were not significant in the sex-stratified PERMANOVA analyses (adjusted *P* > 0.05, **Fig. 3a**), suggesting potential confounding effect of sex on sex-biased microbiome covariates and host phenotype-gut microbiota associations.

**Fig 3.**
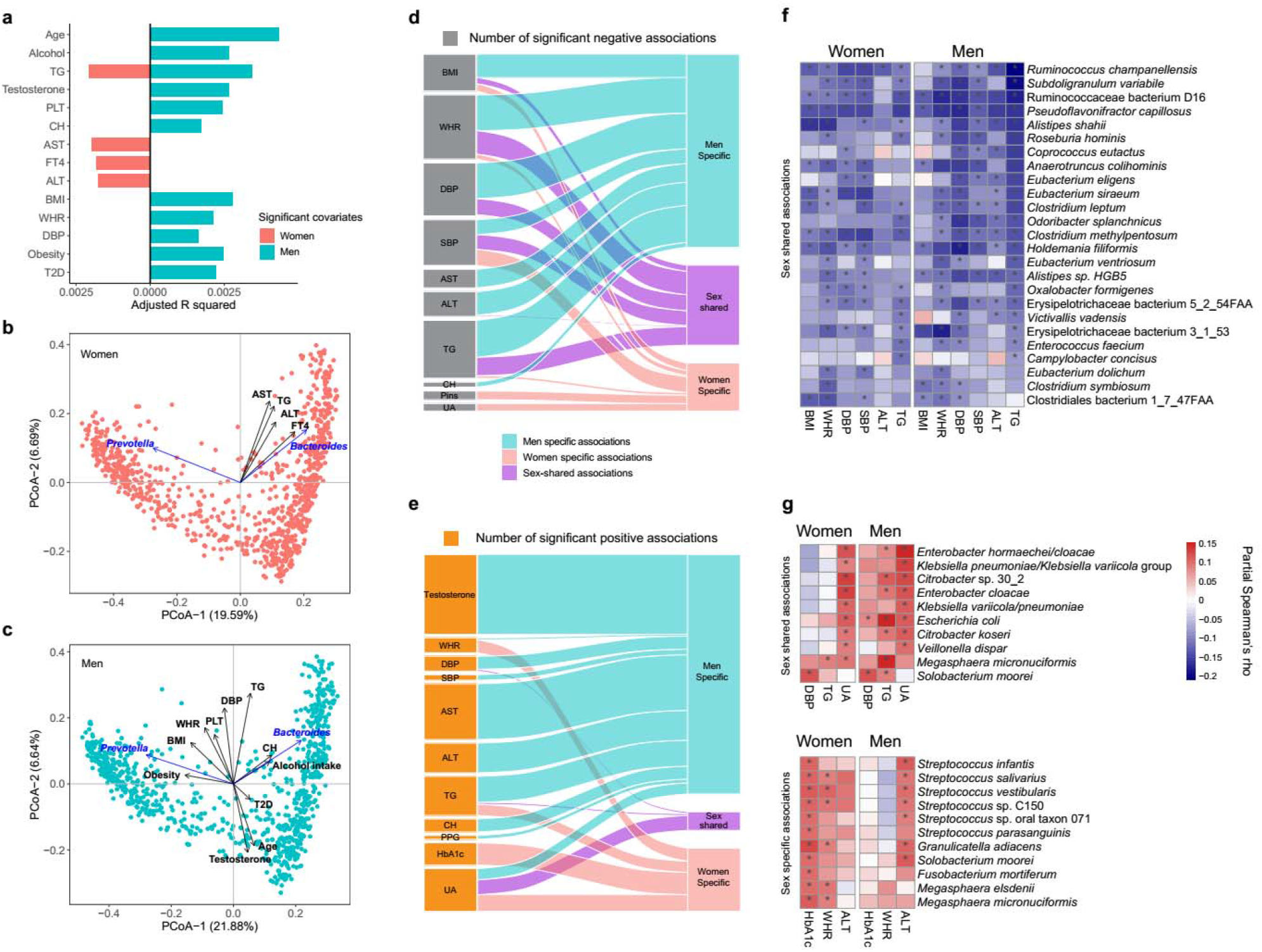
Sex-biased host phenotype-gut microbiota associations. **a,** Horizontal bars showing the amount of inferred variance (adjusted R squared) explained by each identified covariate (adjusted *P <* 0.05) by PERMANOVA using Bray-Curtis dissimilarities at the gene level in women (red) and men (blue). **b-c,** Unconstrained principal coordinate analysis (PCoA) using gene level Bray-Curtis (BC) dissimilarities in women (b) and men (c). Arrows indicate the dimensions of significant covariates as shown in (a) and the contribution of the genera *Bacteroides* and *Prevotella.* **d-e,** Number of significant negative **(d)** and positive **(e)** associations (Partial Spearman’s rank correlations, adjusted *P* < 0.05) between host parameters and individual species specifically in women (red), specifically in men (blue) and shared in both sexes (purple) after adjustment for age. Gray indicates negative associations; orange indicates positive associations. For each panel, the bar height (left) indicates the number of significantly associated species with each phenotype. **f,** Heatmap showing the sex-shared significant negative associations between host parameters and species. **g,** Heatmap showing the sex-shared and selected sex-specific significant positive associations between host parameters and species. See **Extended Data Fig. 5** for full association heatmap. *, adjusted *P* < 0.05.

The gut microbiota of women was characterised by a high degree of overall stability, and unexpectedly, showed no significant associations with age, menopause, and associated metabolic disorders such as obesity, T2D and hypertension (**Fig. 3a-b)**. By contrast, age explained the largest microbial variance in men, and showed consistent direction of the projected impact on the male gut microbiota with testosterone, opposite to those of obesity-related covariates such as BMI, WHR and TG level **(Fig. 3c)**. This finding was in line with the correlations between age, testosterone and obesity-related clinical parameters in men (Spearman’s correlation, adjusted *P* < 0.05, **Extended Data Fig. 4h**).

We next asked whether sex-biased associations between individual species and host parameters could be detected after adjustment for age. A much smaller number of significant associations were identified in women than in men (**Fig. 3d, e**). No significant associations were found between species abundance and sex hormones in women; however, many species were positively correlated with testosterone levels, and largely overlapped with those that were negatively correlated with clinical metabolic parameters in men (Partial Spearman’s correlation, adjusted *P* < 0.05, **Fig. 3e, Extended Data Fig. 5, Supplementary Table 13-14**). Among these species, the abundances of *Faecalibacterium prausnitzii, Roseburia inulinivorans* and *Butyrivibrio crossotus* were persistently and positively correlated with testosterone levels in men after adjustment for both age and BMI (adjusted *P* < 0.05, **Supplementary Table 14**). In agreement with the consistent changes in gut microbial communities in NCDs across the two sexes in previous cross-sectional studies ^9–12, 25^, we observed that several species from the genera *Eubacterium, Alistipes* and *Ruminococcus* were negatively correlated with WHR, blood pressures and TG in both sexes **(Fig. 3d, f)**. Significant positive correlations between the abundance of several Proteobacteria species from the genera *Enterobacter, Citrobacter* and *Klebsiella* and blood UA levels were also shared between the two sexes (**Fig. 3e, g, Supplementary Table 14)**. On the other hand, several *Streptococcus* species were positively correlated with HbA1c and WHR in women, but positively with liver aminotransferase levels rather than any diabetes or obesity-related parameters in men (**Fig. 3e, g, Extended Fig. 5)**. Although several *Streptococcus spp.* have been reported to be enriched in elderly European women with T2D compared to women with normal or impaired glucose control ^33^, further studies are needed to determine the biological role, if any, of these observed sex-biased disease associations.

### Reduced sex microbial differences with ageing

Regardless of sex and ethnicity, ageing is a biological process generally accompanied by impairment of the digestive system and the immune system, and increased multimorbidity and medicine use ^34–37^. However, reported ageing-related gut microbial characteristics are variable and inconsistent ^38^, potentially confounded by ageing-related health conditions, sample size, and sex.

Aiming at extending our knowledge on changes of the gut microbiota in elderly, we compared the gut microbiota of adults below and above 50 years in each sex (**Methods**). In line with the higher impact of age on the overall gut microbiota composition in men, we also observed that the magnitude of differences in microbial α diversity and the proportions of differentially abundant species and functional KOs between younger and elderly individuals were greater in men compared to women **(Fig. 4a-b, Extended Data Fig. 7a, Supplementary Table 15)**. These findings were well replicated in the LLD cohort (**Extended Data Fig. 6a-c, Supplementary Table 15)**. In addition to higher microbial diversity in adults older than 50 years of age, several gut microbial taxonomic and functional characteristics were also shared in elderly adults of both sexes. For instance, comparison of PG adults under and over 50 years of age revealed that PG adults over 50 years of age showed significantly lower B/F ratio and biosynthesis capacity for bacterial lipopolysaccharide (LPS) in gram-negative bacteria and several vitamins, including menaquinone, pantothenate, riboflavin and tetrahydrofolate **(Extended Data Fig. 8, Supplementary Table 15-16)**. Moreover, elderly adults of both the PG and LLD cohorts exhibited higher capacity for microbial methane production as well as a higher abundance of *Methanobrevibacter smithii*, the dominant methanogen in the intestine, than younger adults, and to a greater extent in men than in women **(Extended Data Fig. 8, Supplementary Table 15)**. Given the clinical links established between increased methane production and prolonged intestinal transit time ^39,40^, an age-related increase in methane production might be related to increased chronic constipation in elderly. A second round Permanova analysis further revealed significant impacts of age and obesity-related covariates (obesity, BMI and WHR) on species that differed in abundance between the two age groups in both sexes, but in opposite directions (**Extended Data Fig. 7b-c**). The sex-dependent opposite effects of these obesity-related covariates on the microbiota were consistent with the age- and sex-dependent disparities in metabolic disorders (**Extended Data Fig. 7d, e**). Most blood routine parameters and dietary patterns that differed significantly between the two age groups (**Supplementary Table 3, Extended Data Fig. 7f, g**), showed no significant effects (**Extended Data Fig. 7b**).

**Fig 4.**
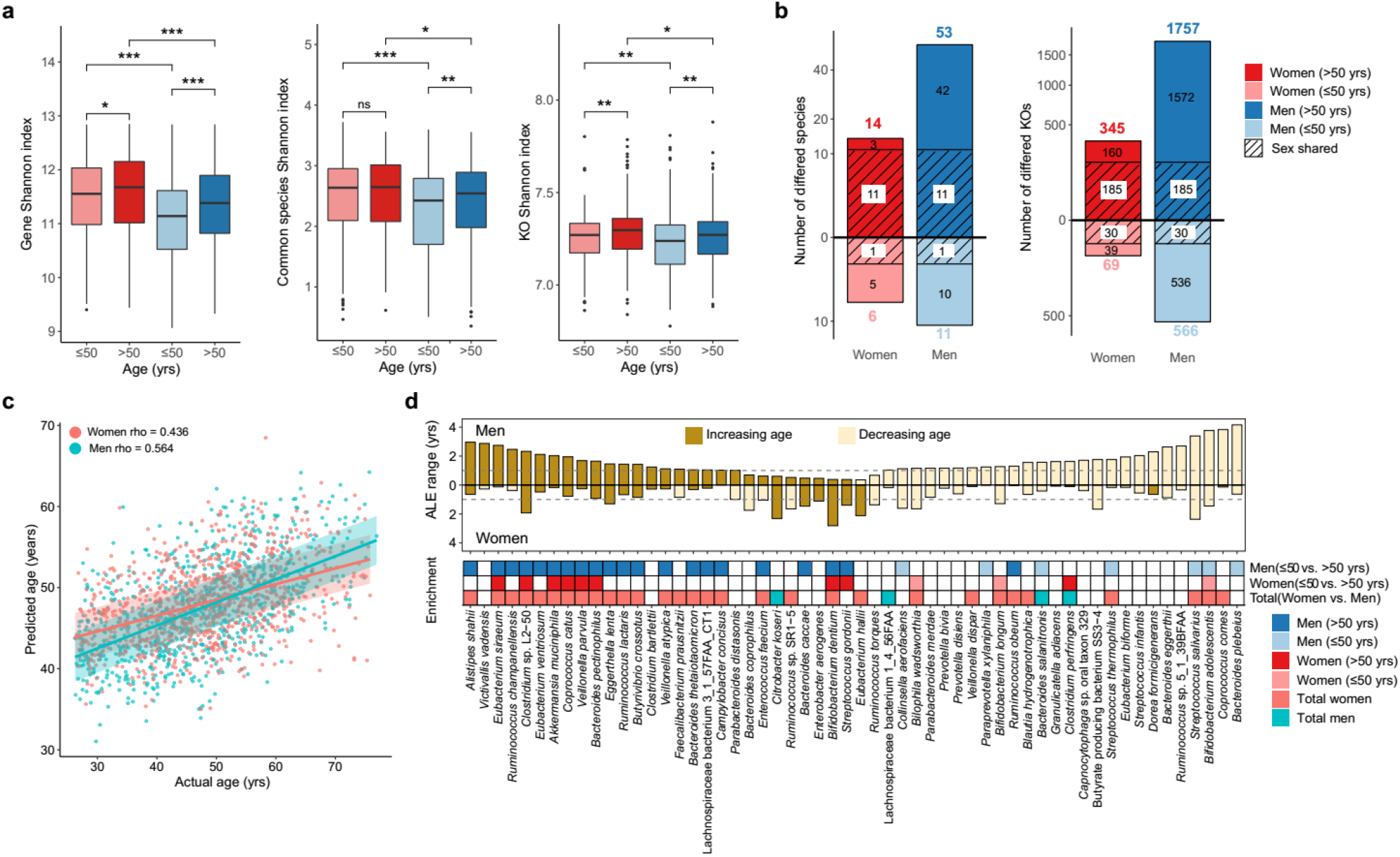
Age- and sex-related differences in the adult gut microbiota of the PG cohort. **a,** Box plots showing gut microbial alpha diversity for adults below (<=50 years, light color) and above 50 years (>50 years, dark color) of women (red) and men (blue) at the gene, common species, and KO level. Wilcoxon rank test; * *P* < 0.05; ** *P* < 0.01; *** *P* < 0.001. **b,** Number of differentially abundant common species (left) and KOs (right) (Wilcoxon rank test, adjusted *P* < 0.05) between two age groups in each sex, with oblique shadow area indicating the fraction shared between the two sexes. **c,** Scatter plot of predicted age versus actual age for women (red) and men (blue) of PG cohort. Spearman’s rho values between predicted age and actual age in each sex are shown. Shaded areas contain 46% for women and 50% for men of predictions corresponding to the trend line ± 3 years. **d,** Accumulated local effects (ALE) range (maximum ALE minus minimum ALE within 5-95% abundance bracket) shown as vertical bars for microbial species affecting age prediction for at least one year, for men (up) and women (down). Dark brown indicates increasing ALEs, light brown indicates decreasing ALEs. The enrichment of each species is shown in the bottom between adults below and above 50 years for men and women, and between all women and all men.

With the purpose of estimating how strong the gut microbiota might be related to host chronological age, we used gut microbial species to build age prediction models in each sex (**Methods**) ^41^. Interestingly, models in both sexes in the PG cohort had good performances, with a relatively higher Spearman’s rho of 0.584 for men compared with 0.436 for women between predicted and actual age **(Fig. 4c)**. Of the models for both sexes, the accumulated local effects (ALE) of most selected features were consistently increasing or decreasing with the predicted ages, but with larger effects in men **(Fig. 4d).** The ALE ranges of selected species features were also sex-biased in the age-prediction models for the LLD cohort **(Extended Data Fig. 6d-e)**. Interestingly, *A. muciniphila, Eubacterium siraeum* and *Coprococcus catus*, were selected as strong positive age predictors only for adult men in both cohorts (ALE range ≥1 year) **(Fig. 4d, Extended Data Fig. 6e**). The elderly enriched species *Streptococcus gordonii* predicted increased average age in both cohorts and was selected as the strongest age predictor for the LLD adults **(Fig. 4d, Extended Data Fig. 6e**). Although the abundances of several *Bifidobacterium* species were decreased in elderly of both cohorts, *B. adolescentis* and *B. longum* were stronger predictors in the PG cohort, whereas *B. bifidum* and *B. animals* more strongly predicted age in the LLD cohort **(Fig. 4d, Extended Data Fig. 6e**).

Additionally, we found significant differences in the abundance of many sex-biased species between age groups. Of note, the abundances of multiple women enriched species were relatively stable among adult women across broad age ranges, but showed significant increases in elderly men as compared to young men **(Fig. 4d**). This raised the question as to whether sex-associated microbial differences were reduced in elderly. In both the PG and the LLD cohorts, sex indeed explained much less of the microbial variances in elderly individuals **(Fig. 5a, c, Supplementary Table 18**), and the number of sex-dependent differences in microbial features was considerably less in elderly compared with younger adults (**Fig. 5b, d**). For instance, 55 and 41 species whose abundance differed between adult women and men of younger age in the PG and the LLD cohort respectively (adjusted *P* < 0.05, **Fig. 5b, d, e**), showed no significant sex difference in elderly ((adjusted *P* > 0.05, **Fig. 5b,d, f**). On the other hand, we found that 38 species differed consitently in abundance between men and women in both age groups in the PG cohort (**Fig. 5f**). Among them, several were correlated with intake of alcohol or smoking in PG men (**Fig. 5f**). For instance, intake of alcohol in men correlated negatively with the abundance of women enriched species, such as *B. adolescentis, R. champanellensis* and *A. shahii*, but positively with men-enriched species *Turicibacter sanguinis* after adjusting for age, BMI, and smoking (**Fig. 5f, Partial Spearman’s correlation, adjusted *P* < 0.05)**. Smoking also negatively correlated with several women-enriched bacteria, including *E. eligens, E. ventriosum, B. crossotus, Haemophilus haemolyticus* and *H. parainfluenzae* **(Fig. 5f)**. Thus, these findings suggest that lifestyle to a certain extent could contribute to the observed sex-dependent differences in the composition of the gut microbiota in the PG cohort.

**Fig 5.**
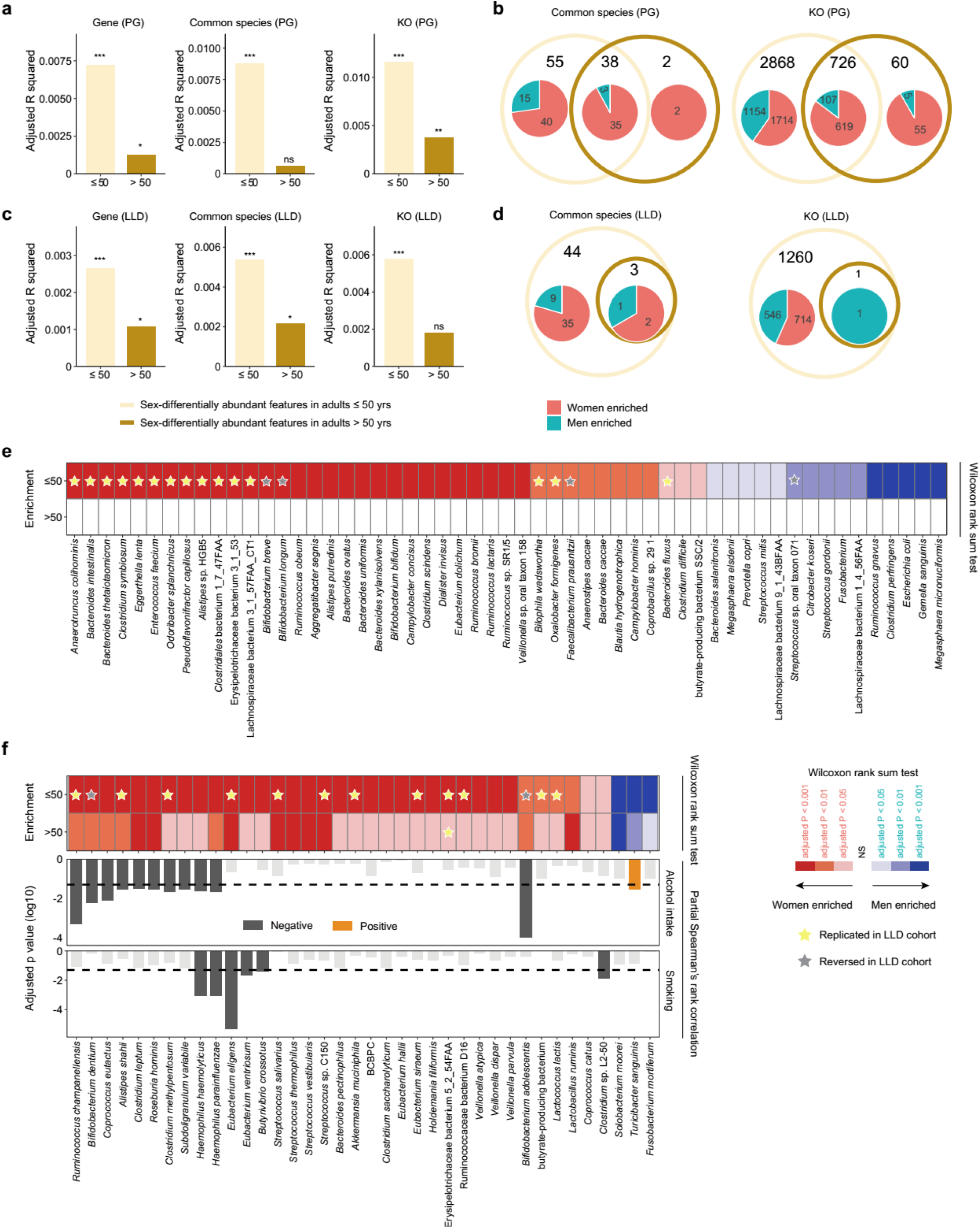
Reduced sex differences in the gut microbiota from young to elderly adults. **a, c,** Sex-explained inferred variance (adjusted R-squared) in the gut microbiota as determined by PERMANOVA with gene (left), common species (middle) and KO (right) level Bray-Curtis dissimilarities for adults below 50 years (light brown) and above 50 years (dark brown) in the PG cohort **(a)** and in the LLD cohort **(c)**. PERMANOVA, * *P* < 0.05, ** *P* < 0.01, *** *P* < 0.001, ns, *P* ≥ 0.05. **b, d,** Venn diagrams showing the number of sex-dependent differentially abundant gut microbial common species (left) and KOs (right) between two age groups of the PG (**b**) and the LLD (**d**) cohort. Pie charts indicate the number of species/KOs enriched in women (red) and men (blue). Wilcoxon rank test, adjusted *P* < 0.05. **e,** Sex-dependent differentially abundant species specifically in adults below 50 years (n=55, panel **b**) in the PG cohort. Stars indicate the enrichment of the respective species replicated (yellow) or reversed (gray) in the LLD cohort (See **Supplementary Table 15** for full list). **f,** Sex-dependent differentially abundant species in adults in two age groups (n=38, panel **b**) in the PG cohort. Additional bar charts (down) showing the significant negative (grey) and positive (orange) associations between these 38 species and alcohol intake or smoking in men after adjusting for age, BMI and one another (Partial Spearman’s rank correlation). Y axis indicates the log (10) transformed adjusted *P* values, with dashed lines indicating adjusted *P* < 0.05 as the cutoff for significance.

## Discussion

Despite emerging evidence linking the gut microbiome to human health, sexual dimorphism in the gut microbiota as well as the potential relation to diseases and behaviour, has often been overlooked. Our study demonstrates the sexual dimorphism in the gut microbiome and associations with age, sex hormones, and other host metadata in a large community-based cohort randomly drawn from a Northern Chinese Han population.

The women in the PG cohort harboured a relatively more diverse and richer gut microbial community than that of men, a finding which was further replicated in published datasets of Chinese and Dutch adults. Of note, both Chinese and Dutch adult women exhibited a significant enrichment of *A. muciniphila,* a well-characterized metabolic beneficial species ^42,43^, and *E. lenta*. The latter species possesses the ability to inactivate digoxin ^44^, the therapeutic effect of which has been reported to differ between women and men ^45^. Significant sex differences were also consistently observed in the abundance of bacterial genes involved in BA transformation and in serum BA profiles, suggesting that the gut microbiota might modulate sex-biased host metabolic processes through BA-Farnesoid X receptor (FXR) and TGR5 signalling pathways. These sex differences of the gut microbiota might potentially be associated with the age-related sex disparities in risk, pathophysiology and treatment of metabolic disorders, where the underlying mechanisms require further investigation.

Focusing on the host-microbiome association patterns in the PG cohort, we show that the overall gut microbial community in women seemed to remain relatively stable between different age groups, while it in men varied significantly with age, testosterone levels, multiple metabolic parameters, and lifestyle. Correspondingly, many species including several butyrate producers were positively correlated with testosterone levels in men but showed negligible relationships with sex hormone levels in women. Additionally, smoking and alcohol intake in men were negatively associated with several women enriched bacterial species, suggesting that the sexual dimorphism of the gut microbiota might be mediated through both hormone-dependent and -independent factors.

Importantly, we demonstrate that the cross-sectional age-related gut microbial differences are much more pronounced in men than in women in the PG cohort, a finding we replicated in the LLD cohort. Despite substantial population and sex differences, commonly shared age-related gut microbial differences were identified, such as increased microbial diversity and methane production potential, and decreased abundances of *Bifidobacterium spp*. in elderly, which might be related to age-dependent changes of intestinal functions. Additionally, we demonstrate considerable accuracy in chronological age prediction from intestinal microbial species abundances in both sexes, again, emphasizing a strong connection between ageing and the human gut microbiota. In contrast to previous reports ^46,47^, we observed that several Firmicutes members from the genera *Eubacterium, Roseburia* and *Clostridium*, exhibited a significant increase in abundance in elderly, especially in elderly men. Among these bacteria, some butyrate producers were independently and positively associated with male testosterone level. As noted, testosterone levels in elderly men in the PG cohort were significantly higher than those in younger men, which is similar to reports of two large-scale cross-sectional Chinese and American cohorts ^48,49^. These findings, which were contrary to an age-related hormone decline, suggest that the decline in serum testosterone in relation to generations is larger than the age-associated decline in cross-sectional population. An intriguing possibility is that social and environmental differences between generations, especially maternal and early life conditions might potentially modify later testosterone levels, gut microbiota, disease susceptibility, and host-microbiome interactions ^50–52^. Further longitudinal studies are needed to clarify the age-dependent changes and generation differences in the gut microbiota that cannot be captured in cross-sectional data, with careful consideration of sex differences.

**Supplementary information** is available online.

## Acknowledgements

This work was supported by grants from National Key Research and Development Program (2016YFC1304901), and Beijing Science and Technology Committee (D131100005313008), and Shenzhen Municipal Government of China (No. JCYJ20170817145809215). We thank Associate Professor Elizabeth K Speliotes from University of Michigan for helpful discussion and suggestions on designing and establishing the Pinggu cohort. We thank Dr. Philipp Rausch from University of Copenhagen and Associate Professor Qiang Sun from University of Toronto for helpful suggestions on statistical methods in the manuscript. We thank the participants and the staff of LifeLines-DEEP for their collaboration. Funding for the project was provided by the Top Institute Food and Nutrition Wageningen grant GH001. The sequencing of the LifeLines-DEEP cohort was carried out in collaboration with the Broad Institute. We gratefully acknowledge colleagues at China National Gene Bank for faecal DNA extraction, library preparation and shotgun sequencing experiments, and helpful discussions. We thank all the participants for agreeing to join this study. We are grateful to the research teams from the endocrinology and metabolic department of Beijing Pinggu Hospital and Peking University People’s hospital for their contribution on field survey and data collection.

## Author contributions

L.J., X.Z. and J.L. designed and coordinated the study. X.Z., Y.L., X. Zhou., and X.H. oversaw the establishment of the Pinggu cohort. Y.L., Z.F. and L.W. were responsible for the overall collection of biological samples and data through field survey. H.Z. led the bioinformatic analyses, and Z.S., Z.Z. and H.R conducted the bioinformatic analyses. S.T., Y. L., D. W., F.Y. and C. F. performed bioinformatic analyses on published phenotypic and metagenomic data of 876 Chinese adults and of 1,135 Dutch adults of the LifeLine DEEP cohort. Z.S., H.Z, Z.Z. and H.R. performed revision of the figures. H.Z. interpreted the data with intensive discussion with X.Z., Z.S., Z.Z., K.K., J.L. and L.J.. H.Z, Z.Z. and X.Z. wrote the first version of the manuscript. H.Z. and K.K. performed substantial revision of the manuscript. All authors participated in discussions and contributed to the revision of the manuscript. All authors read and approved the final manuscript.

## Author information

The authors declare no competing interests.

## Data availability

Metagenomic sequence data of 2,338 faecal DNA samples from the Pinggu cohort have been deposited at China Nucleotide Sequence Archive (CNSA) with the dataset identifier CNP0000381. The published shotgun metagenomic sequence datasets from 876 Chinese adults are available at the NCBI Sequence Read Archive (accession no. SRA045646 and SRA050230) and European Nucleotide Archive (accession no. ERP006678, ERP005860, ERP013562 and ERP013563). The metagenomic sequence data of 1,135 Dutch adults from Lifelines cohort and age and sex information per sample are available at the European Genome-phenome Archive (EGA) with accession no. EGAS00001001704. All other data are available upon request.

## Methods

### Cohort establishment and metadata collection

#### Flowchart for establishment of the Pinggu cohort

The Pinggu cohort study, a large prospective cohort set up in suburban Beijing (in the north of China), was first established in 2013-2014 and designed for follow up studies every 5 years. The Pinggu district is surrounded by mountains on three sides. This unique geographic feature results in a relatively low mobility of the population in the area. The overall research objectives of the Pinggu cohort are to study the involvement of genetic and environmental factors in the development of metabolic diseases and to understand the changes/roles of the gut microbiota on the ageing process.

Based on the national Civil Registration system, a total of 6,583 participants were randomly drawn using multistage stratified sampling method according to the demographic structure in terms of sex, age and region (rural or suburban) **(Fig. 1a)**. Participants were eligible based on the following criteria: 1) born in Pinggu; 2) 5 years or longer continuous residence in Pinggu back tracing from the sampling point, and 3) adults aged 26-76 years (men and non-pregnant women). A total of 4,002 individuals were enrolled, giving a response rate of 60.8%. All enrolled participants signed an informed consent form before their physical examination and biomaterial collection. Subsequently, 2,338 participants meeting the additional criteria: 1) with faecal and blood samples; 2) with complete questionnaire information; 3) without antibiotic treatment in the past 3 weeks before biomaterial collections, and 4) without severe disease (end-stage cancer and renal disease), were selected for the metagenomic study. In total, 98 phenotypic factors were collected for each participants through questionnaires (Q), including socio-demographics (n=5), lifestyle (n=4), diet (n=18), drugs for treating metabolic disorders when collecting faecal samples (n=10), and female gynaecological information (n=4); or through clinical measurements (M), including anthropometric measures (n=4), biochemical measures of blood (n=42) and urine samples (n=1), and liver-to-spleen fat attenuation ratio (L/S ratio) measured using computed tomography (CT) (n=1). Information on diseases (n=9) was collected by self-report of diagnosis and treatment history for patients, and by new diagnoses according to clinical measures for the remaining participants with no self-report. Statistics of the factors are summarized in **Supplementary Table 1**.

Questionnaires, clinical measurements and disease information collected in the study centres are detailed in the following part.

#### Questionnaire

All participants in the Pinggu cohort study provided during a face-to-face interviewer-administered questionnaire information on socio-demographics, medical history, family history of chronic disease, life-style and other health-related topics. The pre-processing of information on sleep duration, sedentary time and diet frequency data was performed as previously described in epidemiological studies on China Kadoorie Biobank comprising 0.5 million Chinese adults ^53,54^. Sleep duration was categorized into 6 or fewer hours, 7-9 h, or more than 9 h ^55^. Sedentary time was categorized into <1.5, 1.5-2.4, 2.5-3.4, 3.5-4.4, or ≥ 4.5 hours/day ^56^. The average frequency of consumption of food items was categorized into ‘never/rarely’, ‘monthly’, ‘1-3 days per week’, ‘4-6 days per week’, and ‘daily’ ^57^.

#### Physical Examination

Clinical measurements, height, weight, waist and hip circumferences were measured by trained staff according to standardized protocols. The derived parameters body mass index (BMI, kg/m2) and waist-to-hip ratio (WHR) were calculated and stored for further analyses. Blood pressure (mmHg) was measured three times using an automatic manometer (Omron, Japan) after a 10-min period of rest in a seated position and the mean of the measures was used in the analysis.

#### Laboratory measurements and biobanking

All participants had fasting blood samples drawn between 8:00 AM and 9:00 AM for the assessment of fasting levels of glucose, insulin, lipids, sex hormones and other relevant biochemicals. Afterwards, a standard oral glucose tolerance test (OGTT) was conducted for participants with no known diabetes. Blood samples were then drawn for the assessment of 2-hour postprandial glucose and insulin levels. Urine samples were collected in fasting conditions in the visit morning, and faecal samples were collected using sterile cups after defecation at visit. Details for laboratory measurements are summarized in **Supplementary Table 19**. Portions of these biomaterials including blood, urine and faecal samples were also stored at −80 °C for long-term biobanking in addition to current measurement use.

### Definition of diseases or conditions

#### Definition of obesity

Participants were classified as normal weight (BMI < 24 kg/ m^2^), overweight (24.0 ≤ BMI < 28 kg/m^2^), or obese (BMI ≥ 28 kg/m^2^) according to criteria issued by the China Diabetes Society ^58^.

#### Definition of hypertension

Hypertension was defined using blood pressure of at least 140/90 mmHg or the current use of antihypertensive medications.

#### Definition of prediabetes and diabetes

Known type 2 diabetes was defined by a self-reported history of diabetes diagnosed by a doctor and/or on glucose lowering treatment. Participants without known diabetes underwent a 75g 2-h oral glucose tolerance test (OGTT). According to the WHO definition in 1999 ^59^, undiagnosed diabetes was defined as fasting plasma glucose (FPG) ≥7.0mmol/L and/or 2-h postprandial plasma glucose (PPG) ≥11.1mmol/L. Prediabetes was defined as 6.1 ≤ FPG <7.0 mmol/L or 7.8 ≤ PPG < 11.1 mmol/L. Normal glucose tolerance (NGT) was defined as FPG <6.1 mmol/l and PPG < 7.8mmol/L.

#### Definition of dyslipidaemia

Dyslipidaemia was defined as CH > 200 mg/dL (5.18 mmol/L), and/or TG>150 mg/dL (1.70 mmol/L), and/or LDL>130mg/dL (3.37 mmol/L), and/or HDL< 40 mg/dL (1.04 mmol/L) or the current use of anti-dyslipidaemia medications ^60^.

#### Definition of hyperuricemia

Hyperuricemia was defined by a serum UA concentration > 416.4 μmol/l (7.0 mg/dl) in men or >356.9μmol/l (6.0mg/dl) in women or a history of gout ^61^.

#### Definition of fatty liver disease

To detect fatty liver disease, unenhanced abdominal CT scans were run for each participant using a 64-slice multi-detector scanner (LightSpeed VCT, General Electric Healthcare, Milwaukee, WI, USA). The Hounsfield Units (HU) of three 1cm^2^ areas in liver and two 1cm^2^ areas in spleen were measured. The mean of the liver and spleen measurements were used to calculate the L/S ratios. Fatty liver disease (FLD) was then defined as : 1) a negative test of HBsAg and anti-HCV, 2) no other special cause of secondary hepatic disease, 3) L/S ratio≤ 1.1 ^23^, and based on 4) history of significant alcohol consumption [men >210g/week or women >140g/week] further classified as alcoholic fatty liver disease (AFLD), or else non-alcoholic fatty liver disease (NAFLD).

## Methods for Metagenomics

### 1. Generation and profiling of shotgun metagenomic sequencing data of Pinggu cohort

DNA extraction from faecal samples was performed as previously described ^9^. DNA nanoball (DNB) based DNA library construction and combinatorial probe-anchor synthesis (cPAS) based shotgun metagenomic sequencing with 100bp single-end reads were applied to all 2,338 samples (MGI, Shenzhen, China). Quality control (QC) workflow developed for this platform was applied to filter out low-quality and human reads ^62^. On average 6.9 Gb (± 2.1 Gb) high-quality data was generated per sample. High-quality non-human reads were further aligned to the 9.9M integrated gene catalogue (IGC) ^28^. To control for the quantitative biases of fluctuations in sequencing depth, the IGC uniquely mapped reads were downsized to 20 million for each sample and then used to generate the relative abundance profiles of genes, phyla, genera, species and KOs per individual. A total of 26 phyla, 316 genera, 525 species and 6865 KOs were detected. At species level, we further confined our analyses to species with at least 100 annotated genes in each of at least 10% samples. This gave 151 common species accounted for on average 99.45% of the annotated microbial species composition.

### 2. Richness and diversity analyses

Alpha diversity quantified by the Shannon index was calculated on the relative abundance profiles at gene, KO, and species level using the function *diversity* in the R package *vegan* (R version 3.5.1). Richness was derived as the count number of genes and KOs in each sample as described ^28^.

### 3. Available shotgun metagenomic datasets from Chinese and Dutch adults

To provide a landscape of Chinese adults’ gut microbiota, we further retrieved metagenomic datasets of 876 Chinese adults (aged 18-86 years) from five published studies ^9–12, 25^, in which, samples from patients with diseases known to exhibit severe dysbiosis of the gut microbiota, including liver cirrhosis and extreme obesity (BMI > 32 kg/m^2^) were excluded. Of all published shotgun metagenomic datasets, the Lifelines DEEP (LLD) study from Netherland has a comparable cohort size and age range (1,135 Dutch aged 18-80 years) as the PG cohort ^27^ and was thus retrieved with sex and age information for validation purpose of sex and age-related gut microbial differences. Considering that the sequencing depth was considerably lower in the published Chinese datasets and the LLD cohort ^27^ (3 Gb on average) compared to the PG cohort (6.9 Gb on average), these shotgun metagenomic sequence data were analyzed using the same IGC-based pipeline as described above but without downsizing. The summary of metagenomic datasets is provided in **Supplementary Table 20**.

### 4. Enterotyping for Chinese and Dutch adults

Due to a relatively high gut microbial abundance of *Bifidobacterium* in the LLD cohort, the Dutch metagenomic data could not fit a recently established standard classifier for enterotyping ^63^. Following the guidelines in the aforementioned paper, *de-novo* genus-level enterotyping was performed respectively for 3,214 Chinese adults (2,338 from the PG cohort and 876 from the published dataset), and for 1,135 LLD Dutch adults, according to the partitioning around medoids (PAM) clustering approaches based on Jensen-Shannon divergence (PAM-JSD) or Bray-Curtis dissimilarity (PAM-BC) from Arumugam et al ^64^. The optimal number of enterotype (ET) clusters was evaluated using the Calinski–Harabasz index, which indicated two optimal clusters (ET-*Bacteroides* and ET-*Prevotella*) for Chinese adults and three optimal clusters (ET-*Bacteroides*, ET-Firmicutes and ET-*Bifidobacterium*) for LLD Dutch adults. We further tested the robustness of the optimal number of enterotypes in Chinese and Dutch adults by repeating PAM-BC and PAM-JSD enterotyping on randomly sampled subsets with size ranging from 200 to the maximal number available from each cohort increasing by 100 samples each time. For each given subset size, the procedure was repeated 100 times. For Chinese, the optimal number is consistently 2 in over 97.4% of the test sets. For Dutch adults, the optimal number of enterotype cluster shifts to three as the sample size increased.

### Method for serum bile acid measurements

Serum bile acids in 424 individuals of a selected age-, sex- and BMI-matched sub cohort were measured using the procedure adopted from Sun *et al* ^65^, with minor modifications. Briefly, the bile acid concentrations were measures using Ekspert ultraLC-100 coupled to a Triple TOF 5600 system (AB SCIEX). Fourteen bile acid standards, including cholic acid (CA), ursodeoxycholic acid (UDCA), hyodeoxycholic acid (HDCA), chenodeoxycholic acid (CDCA), deoxycholic acid (DCA), taurocholic acid (TCA), tauroursodeoxycholic acid (TUDCA), taurohyodeoxycholic acid (THDCA), taurochenodeoxycholic acid (TCDCA), taurodeoxycholic acid (TDCA), glycocholic acid (GCA), glycoursodeoxycholic acid (GUDCA), glycochenodeoxycholic acid (GCDCA), glycodeoxycholic acid (GDCA) and internal standard chlorpropamide were purchased from Sigma-Aldrich. Chromatographic separation was achieved on an XBridge Peptide BEH C18 column (100 mm × 2.1 mm i.d., 1.7 μm, Waters Corp.). Column temperature was 40 °C, and the flow rate was 0.4 ml/min. The mobile phase included a mixture of 0.1% formic acid and 10 mM acetamide in water and 0.1% formic acid and 20% acetonitrile in methanol.

## Statistical analyses

### 1. Impacts of drugs

Of the 2,338 individuals, 597 (25.5%) were taking at least one type of drugs for treating diabetes, hypertension or dyslipidaemia around the time for donating faecal sample (**Supplementary Table 2**). Of these drugs registered in the PG cohort, ten with at least 20 users, including antidiabetic drugs (biguanides, glucosidase inhibitors (GIs), sulphonylureas and glinides), antihypertensive drugs (calcium channel blockers (CCBs), beta-adrenergic receptor blockers (beta blockers), diuretics, angiotensin II receptor blockers (ARBs), angiotensin-converting enzyme inhibitors (ACEIs)), and statins for dyslipidaemia, were used for analyses.

To evaluate the impacts of each type of the ten drugs on the gut microbiota, we conducted multiple rounds of PERMANOVA analyses based on Bray-Curtis dissimilarities at the gene, species, and KO level. PERMANOVA analyses were conducted using the function *adonis* from the *vegan* R package. R-squared (R^2^) was adjusted for the number of observations and the number of degrees of freedom using the function *RsquareAdj* from the same package. The *P value* was determined by 10,000 permutations and was further adjusted for multiple testing of tested drugs in each round using Benjamini-Hochberg (BH) method (function *p.adjust,* package *stats*) ^66^. An adjusted *P* value smaller than 0.05 was considered statistically significant. For each round, 1,741 participants not using any of the registered drugs were used as treatment naive controls. In the first round, for a given drug, PERMANOVA was performed on datasets from participants taking it alone or in combination with others and treatment native controls. Drugs such as biguanides, GIs, glinides, sulphonylureas, ARBs and statins were identified to be of significance (adjusted *P* < 0.05, **Supplementary Table 2**). In the next round, samples from participants taking biguanides and/or GIs, which showed the most significant impacts on the overall gut microbial variations in the first round, were excluded. The effect remained significant for ARBs and statins, but not for glinides and sulphonylureas. In the last round, PERMANOVA was performed on samples from individuals taking only one type of drug and treatment naive controls to evaluate the standalone effect of the drug. Of the drugs that significantly influenced the composition of the gut microbiota identified above, GIs were not evaluated in this round due to a shortage of samples, and no other drugs than biguanides were still found to significantly impact on the composition of the gut microbiota (adjusted *P* < 0.05, **Supplementary Table 2**). Taken together, these results demonstrate the complex *in vivo* effects of drugs on the gut microbiota, especially when taken in combination, implying great caution for evaluation and interpretation of result from cohort studies. Subsequent analyses if not otherwise indicated were performed on the 1,741 individuals with no registration of intake of drugs, denoted as ‘the analysis cohort’.

### 2. Association analysis between phenotypic factors and microbial Bray-Curtis dissimilarity

To examine the association between host phenotypic factors and the gut microbiota, PERMANOVA was performed on the entire analysis cohort and further on each sex of the analysis cohort to assess whether there were different association patterns between the two sexes. PERMANOVA was conducted as described above and threshold for statistical significance was BH adjusted *P* values below 0.05.

Bray-Curtis dissimilarities at the gene level were visualized by unconstrained principal coordinate analysis (PCoA) plots with arrows indicating the dimensions of top strong covariates identified by PERMANOVA and the contributions of the genera *Bacteroides* and *Prevotella* were fitted to the ordination space using maximum correlation (*envfit* function, R package *vegan*).

### 3. Principal component analysis for metadata

Principal component analysis (PCA) was implemented using the R function *prcomp* on three subsets of metadata: blood routine parameters, metabolic parameters, and dietary patterns for the entire analysis cohort, women only and men only. Before PCA, all factors were transformed using log transformation. A total of 18 metabolic parameters associated with metabolic disorders including obesity (BMI and WHR), hypertension (SBP and DBP), diabetes (HbA1c, FPG, PPG, fasting insulin [Fins], postprandial insulin [Pins] and HOMA-IR), dyslipidaemia (TG, CH, HDL and LDL) and hyperuricemia (UA), and fatty liver disease (L/S ratio, ALT and AST) were included for PCA analysis. For full information of factors of blood routine parameters (n=16) and dietary patterns (n=18) see **Supplementary Table 3**.

### 4. Age Grouping

In the PG cohort (n=2,338), 50 years was both the median menopause age for women and the median age for men (**Supplementary Table 1**), in agreement with large-scale epidemiological studies in China ^67^ and Europe ^68^ the reported prevalence of multimorbidity increases substantially from this age. We have thus divided both women and men into two groups by an age cut-off of 50 years: a younger group (26 < age ≤ 50 years) and an elderly group (50 < age ≤ 76 years).

### 5. Comparisons on phenotypic factors and microbial characteristics across sexes and age groups

Wilcoxon rank sum tests were applied to detect differences in the continuous metadata and gut microbial features (richness, diversity, relative abundances of phyla, species and KOs) between groups. Chi-square tests were conducted to detect differences in categorical metadata. BH adjusted *P* value less than 0.05 was considered significant.

Differentially enriched KEGG pathways (modules) between groups were identified according to the reporter Z-scores of all KOs involved in a given pathway (module) ^69^. An absolute value of reporter score ≥ 1.96 (95% confidence according to normal distribution) was used as the detection threshold for significance. Of note, in comparative analysis of female age groups, we have excluded 61 postmenopausal women in the younger group and 37 premenopausal women in the elderly group.

### 6. Chronological age prediction from gut microbial species

To test how strong gut microbial features related to age, an age predictor for each sex was trained based on the relative abundance of microbial species with at least 500 represented genes. The predictors were trained as a regressor with five-fold cross-validation using XGboost from *R* package *caret* as recently reported ^41^. After completing grid search for various model configurations, the best performing model was selected based on the minimal RMSE (Root Mean Square Error). For PG adults, the best performing XGBoost model for women was derived with the following parameters: nroduns = 1500, eta = 0.01, max_depth = 2, gamma = 0.9, colsample_bytree = 1, min_child_weight = 2, subsample = 0.5. The best performing XGBoost model for men was derived with the following parameters: nroduns = 5000, eta = 0.01, max_depth = 4, gamma = 0, colsample_bytree = 0.4, min_child_weight = 3, subsample = 0.5. For LLD adults, the best performing XGBoost model for women was derived with the following parameters: nroduns = 1300, eta = 0.01, max_depth = 4, gamma = 0.5, colsample_bytree = 0.8, min_child_weight = 2, subsample = 0.5. The best performing XGBoost model for men was derived with the following parameters: nroduns = 3100, eta = 0.005, max_depth = 2, gamma = 0.1, colsample_bytree = 0.4, min_child_weight = 3, subsample = 0.75.

### 7. Association analyses in the PG cohort

#### Association analyses between phenotypic factors in the PG cohort

Spearman’s rank correlation (SCC) analysis was performed to detect associations between phenotypic factors including age, sex hormones, and metabolic parameters.

#### Association analyses between serum bile acids and gut microbial features in the PG cohort

SCC analysis was performed to detect associations between relative concentrations of serum bile acids and abundances of microbial species / microbial BA transformation genes in 424 selected samples.

#### Association analyses between phenotypic factors and gut microbial features in the PG cohort

A first round of Spearman’s rank correlation (SCC) analysis was performed to detect interactions between host factors and microbial features (microbial diversity, richness and species abundance) without controlling for the potentially confounding effects from other microbial covariates **(Supplementary Table 9, 12)**. Due to the sex disparity in associations between age, metabolic parameters, and overall gut microbial variation, partial Spearman’s rank correlation analyses were further conducted to validate the SCC identified associations between phenotypic factors (sex hormones and 18 metabolic parameters) and microbial features by adjusting for age, or both age and BMI **(Supplementary Table 10-11, 13-14)**. For associations between alcohol intake and microbial features in men, partial Spearman’s rank correlation analysis was conducted by adjusting for age, BMI and smoking, and for associations between smoking and microbial features, age, BMI and alcohol intake were adjusted. The *P* values were adjusted using BH method for total number of tests for each phenotypic factor and the significant cut-off was set at BH adjusted *P* < 0.05.

