## Supplemental Data 1 for "Age-dependent sexual dimorphism in the adult human gut microbiota"

**Supplementary information of “*Age-dependent sexual dimorphism in the adult human gut microbiot*a”**

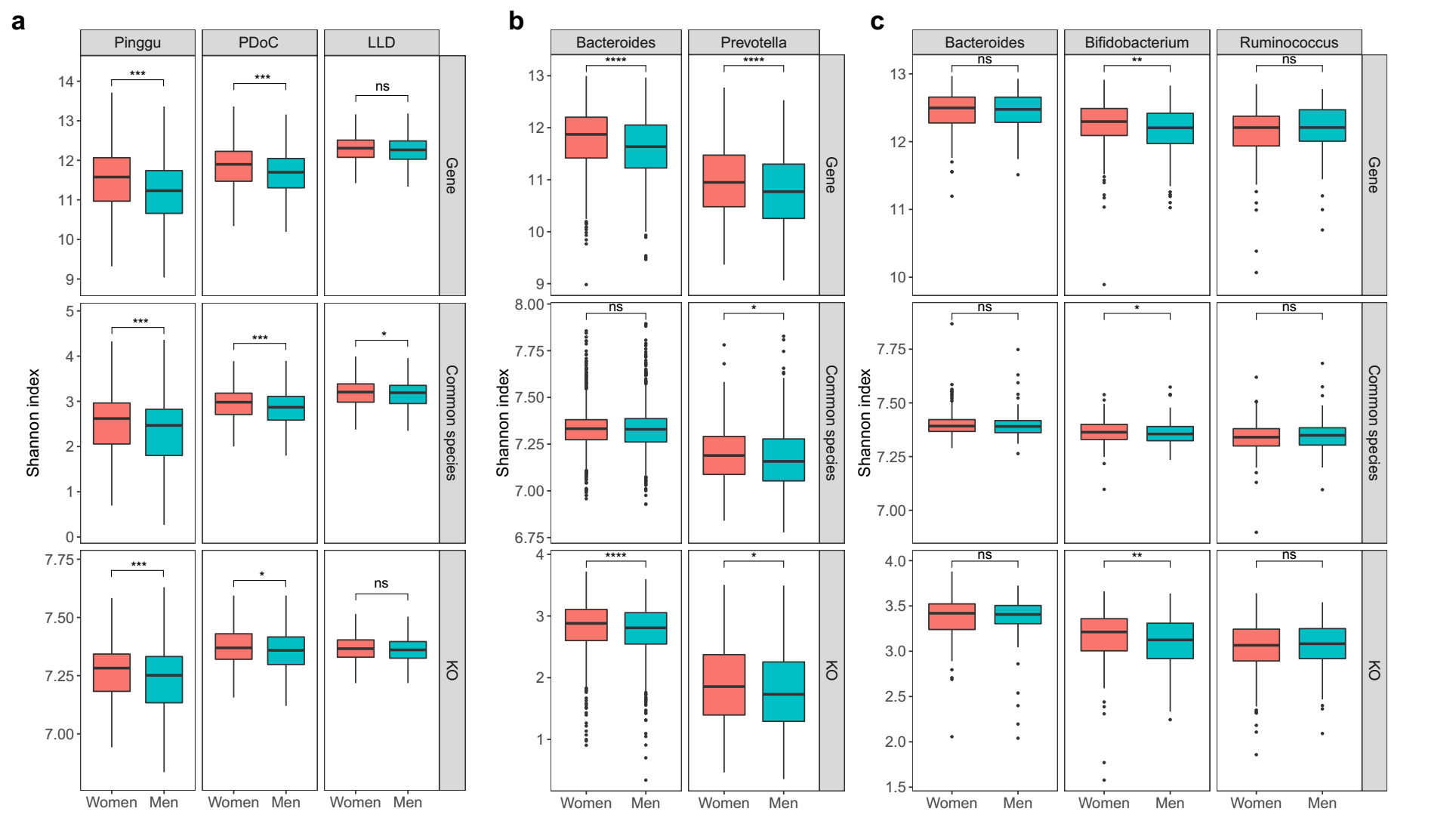

**Extended Data Fig. 1. Sex differences in gut microbial diversity**

**a,** Box plots showing gut microbial alpha diversity between women (red) and men (blue) based on shotgun metagenomic data for the Pinggu (PG, n=2,338), the published datasets of Chinese (PDoC, n=876) and the LifeLines DEEP (LLD, n=1,135) adults at the gene (top), common species (middle), and KO (bottom) level.

**b,** Box plots showing gut microbial alpha diversity between two sexes in the two enterotypes of Chinese adults (n=3,214, including 2,338 from the PG cohort and 876 from published datasets).

**c,** Box plots showing gut microbial alpha diversity between two sexes in the three enterotypes of LLD Dutch adults. Wilcoxon rank test; * *P*  < 0.05; ** *P* < 0.01; *** *P* < 0.001.

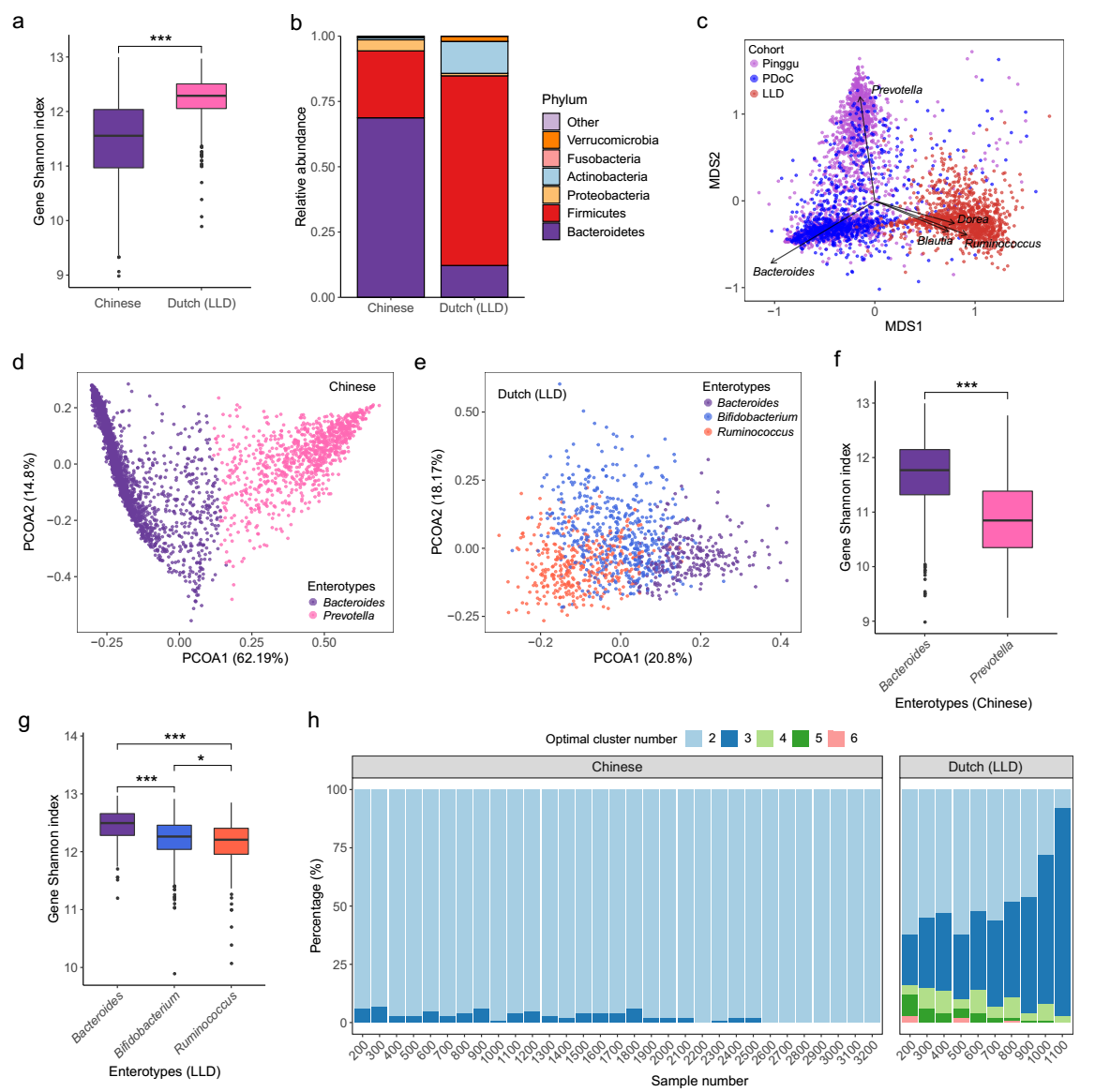

**Extended Data Fig. 2. Comparison of the gut microbial composition between Chinese and Dutch adults.**

**a,** Box plot showing the gut microbial alpha diversity of Chinese (n=3,214) and LLD Dutch (n=1,135) adults. Wilcoxon rank test, *** *P* < 0.001.

**b,** Bar chart showing the mean relative abundance of each bacterial phylum in Chinese (left) and Dutch adults (right).

**c,** Unconstrained Principal coordinates analysis (PCoA) plotting Bray-Curtis (BC) dissimilarities based on the genus profile of Chinese and Dutch adults. Top 5 contributing genera are displayed with arrow length scaled according to the correlation between the genus and PCoA ordination dimensions.

**d,** Scatter plot indicating the two optimal enterotypes identified in Chinese adults, driven by *Bacteroides* (purple) and *Prevotella* (pink), using the BC-based Partitioning Around Medoids (PAM) clustering method.

**e,** Scatter plot indicating the three optimal enterotypes identified in Dutch adults, driven by *Bacteroides* (purple), *Ruminococcus* (orange) and *Bifidobacterium* (light blue).

**f-g,** Box plot showing alpha diversity of the two enterotypes of Chinese adults (**f**) and of the three enterotypes of LLD adults (**g**). Wilcoxon rank test, * *P* < 0.05, *** *P* < 0.001.

**h,** Optimal number of enterotypes identified in 100 times of randomly sampled subsets in metagenomic datasets of Chinese and LLD adults, respectively. Sample number on the x-axis represents the sample size of each subset.

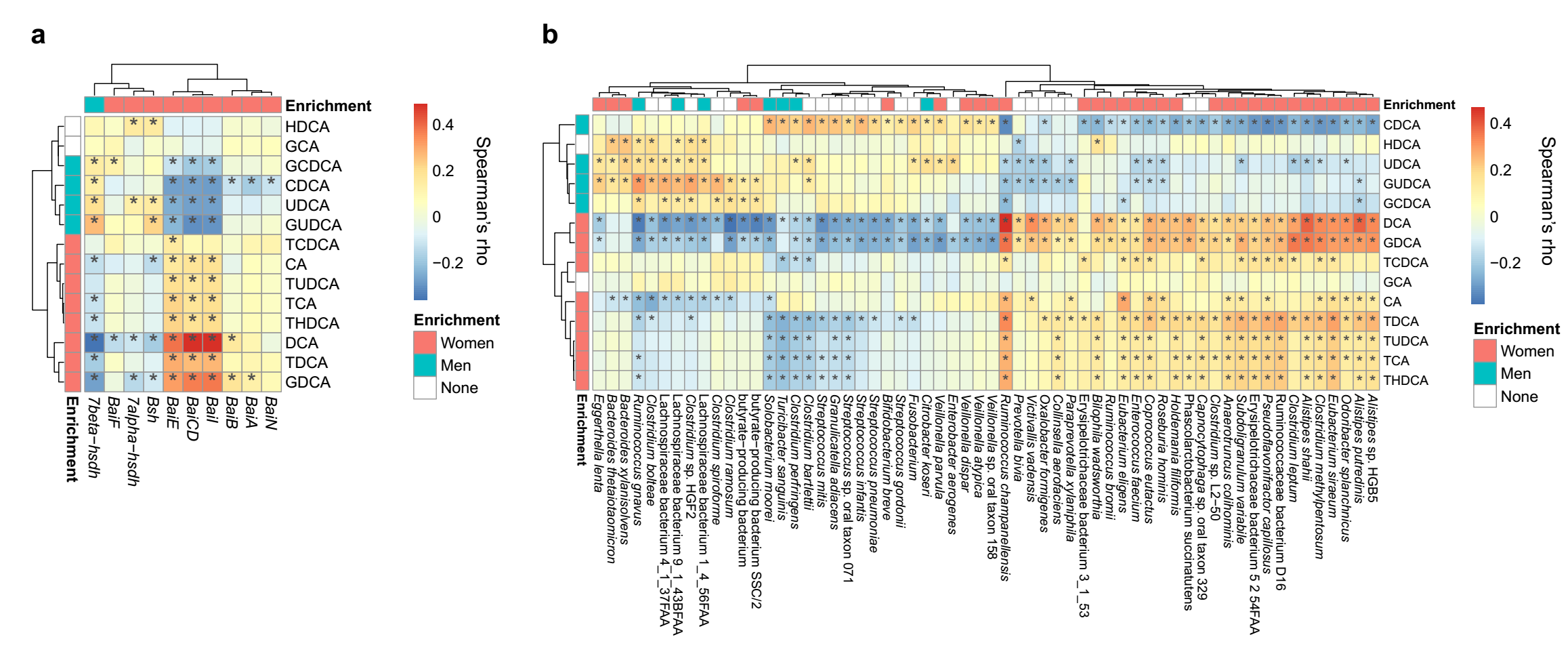

**Extended Data Fig. 3. Associations between serum bile acid and gut microbial genes/species**

**a,** Heatmap showing Spearman’s rank correlation between relative concentrations of serum bile acids and relative abundance of genes involved in bile acid transformation and transport in a subgroup including 424 age-, sex-, and BMI-matched PG individuals. *, adjusted *P* < 0.05.

**b,** Heatmap showing Spearman’s rank correlation between relative concentrations of serum bile acids and relative abundance of common species. *, adjusted *P* < 0.05. Only associations with absolute values of Spearman’s Rho ≥0.2 are shown.

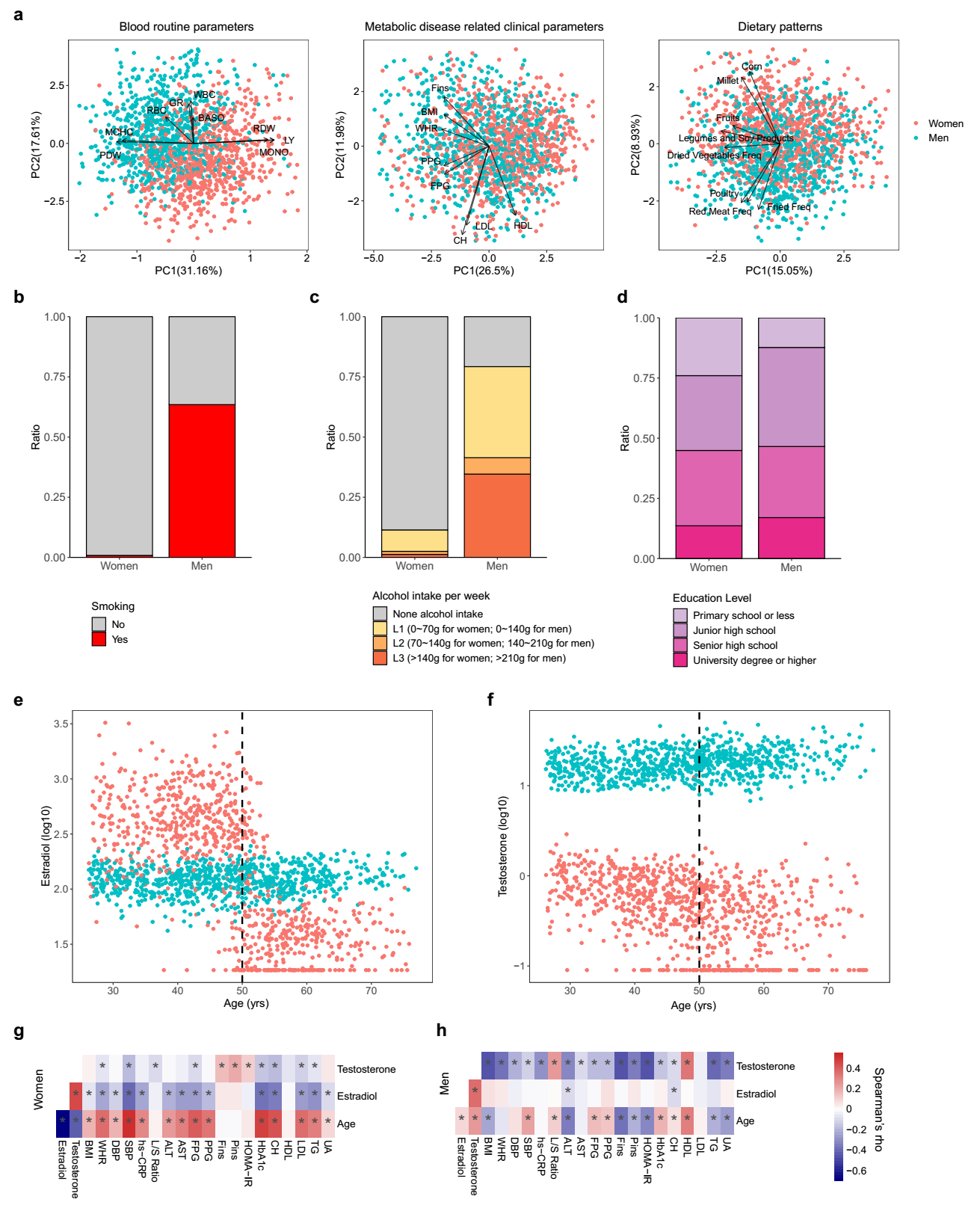

**Extended Data Fig. 4. Sex differences in clinical parameters and lifestyle.**

**a,** Principal components analysis (PCA) of blood routine parameters (left), of metabolic parameters (middle) and of dietary patterns measured/collected in women (red) and men (blue). For each panel, factors with absolute component scores for PC1 or PC2 ≥ 0.3 are shown as primary contributors.

**b-d,** Bar plot showing the distribution of smoking **(b)**, averaged intake amount of alcohol per week **(c)** and education level **(d)** among individuals of each sex.

**e-f,** Serum levels of estradiol **(e)** and testosterone **(f)** by age in women (red) and men (blue), with dashed line indicating the median age 50 years of menopause for the PG women.

**g-h,** Spearman’ s rank correlation between selected host factors in all women (g) and in all men (h). *, adjusted *P* *<* 0.05.

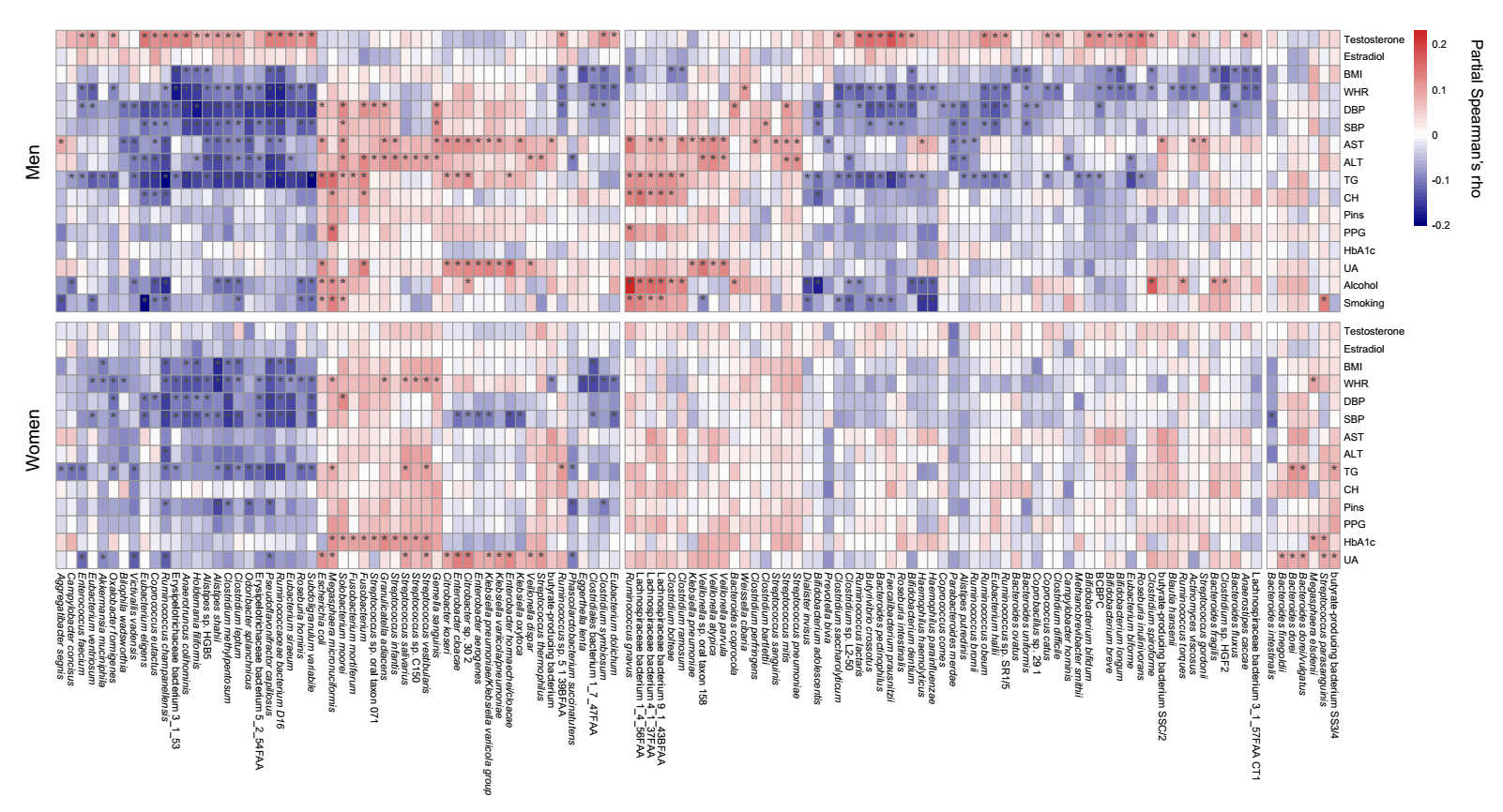

**Extended Data Fig. 5. Correlations between selected host phenotypes and gut microbial species.**

Heatmap showing significant correlations between selected host phenotypes and species abundance in men (up) and in women (down) after adjustment for age. Partial Spearman's correlation. *, adjusted *P* < 0.05.

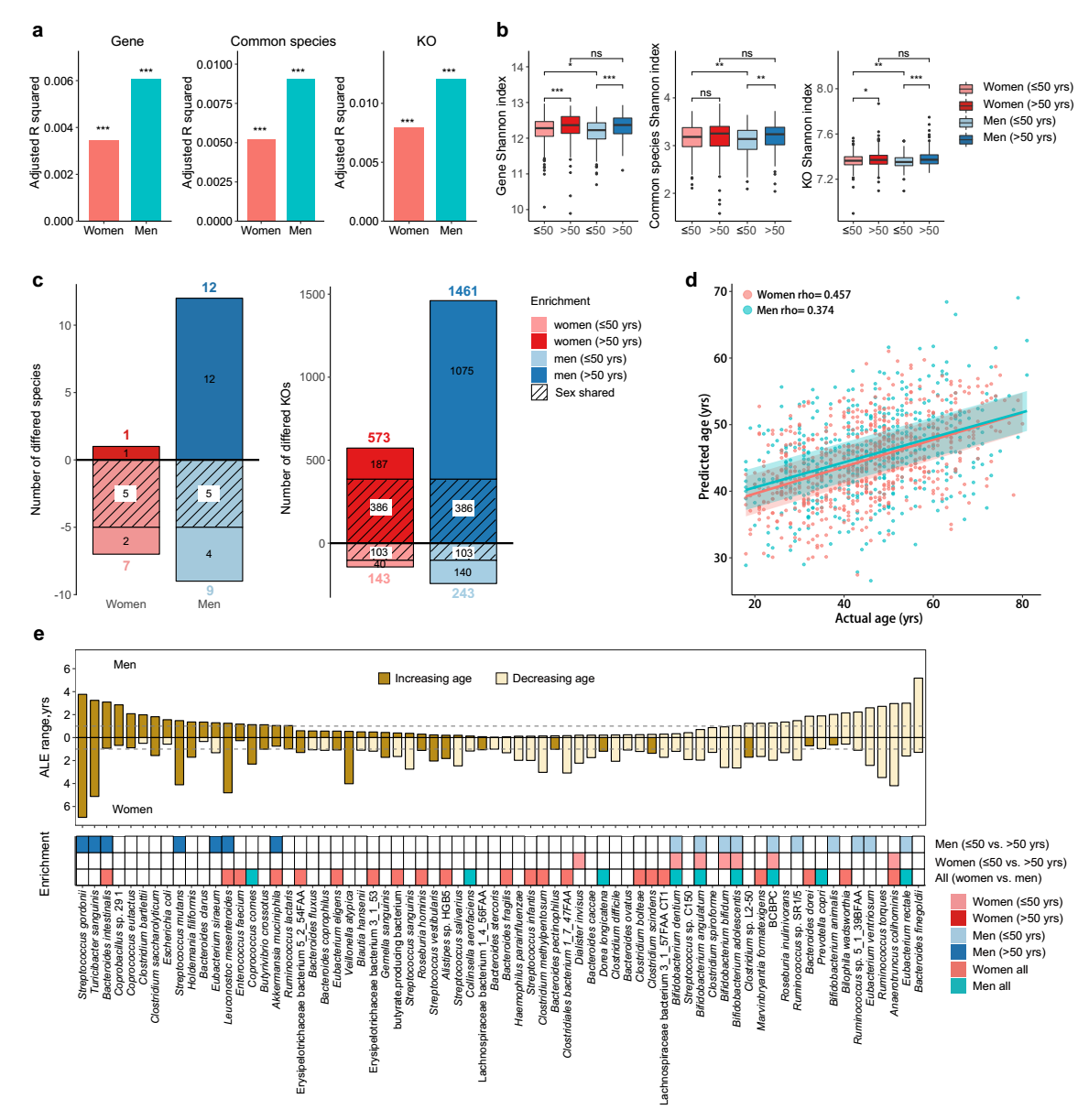

**Extended Data Fig. 6. Age- and sex- related differences in the adult gut microbiota of the LLD cohort.**

**a,** Age-explained inferred variance (adjusted R-squared) in gut microbiome as determined by PERMANOVA with gene (left), common species (middle) and KO (right) level Bray-Curtis dissimilarities for women (red) and men (blue).

**b,** Box plots showing gut microbial alpha diversity for women (red) and men (blue) below (<=50 years, light color) and above 50 years (>50 years, dark color) at the gene, common species and KO level. Wilcoxon rank test; * *P* < 0.05; ** *P* < 0.01; *** *P* < 0.001.

**c,** Number of differentially abundant common species (left) and KOs (right) (Wilcoxon rank test, adjusted *P* < 0.05) between the two age groups in each sex, with oblique shadow area indicating the fraction shared between the two sexes.

**d**, Scatter plot of predicted age versus actual age for women (red) and men (blue) of PG cohort, with Spearman's rho indicated on top. Shaded areas contain 46% for women and 50% for men of predictions and correspond to the trend line ± 3 years.

**e**, Accumulated local effects (ALE) range (maximum ALE minus minimum ALE within 5-95% abundance bracket) shown as horizontal bars for microbial species affecting age prediction by at least 1 year, for men (up) and women (down). Dark brown indicates increasing ALEs, light brown indicates decreasing ALEs. The enrichment of each species is shown in the bottom between adults below and above 50 years for men and women, and between all women and all men.

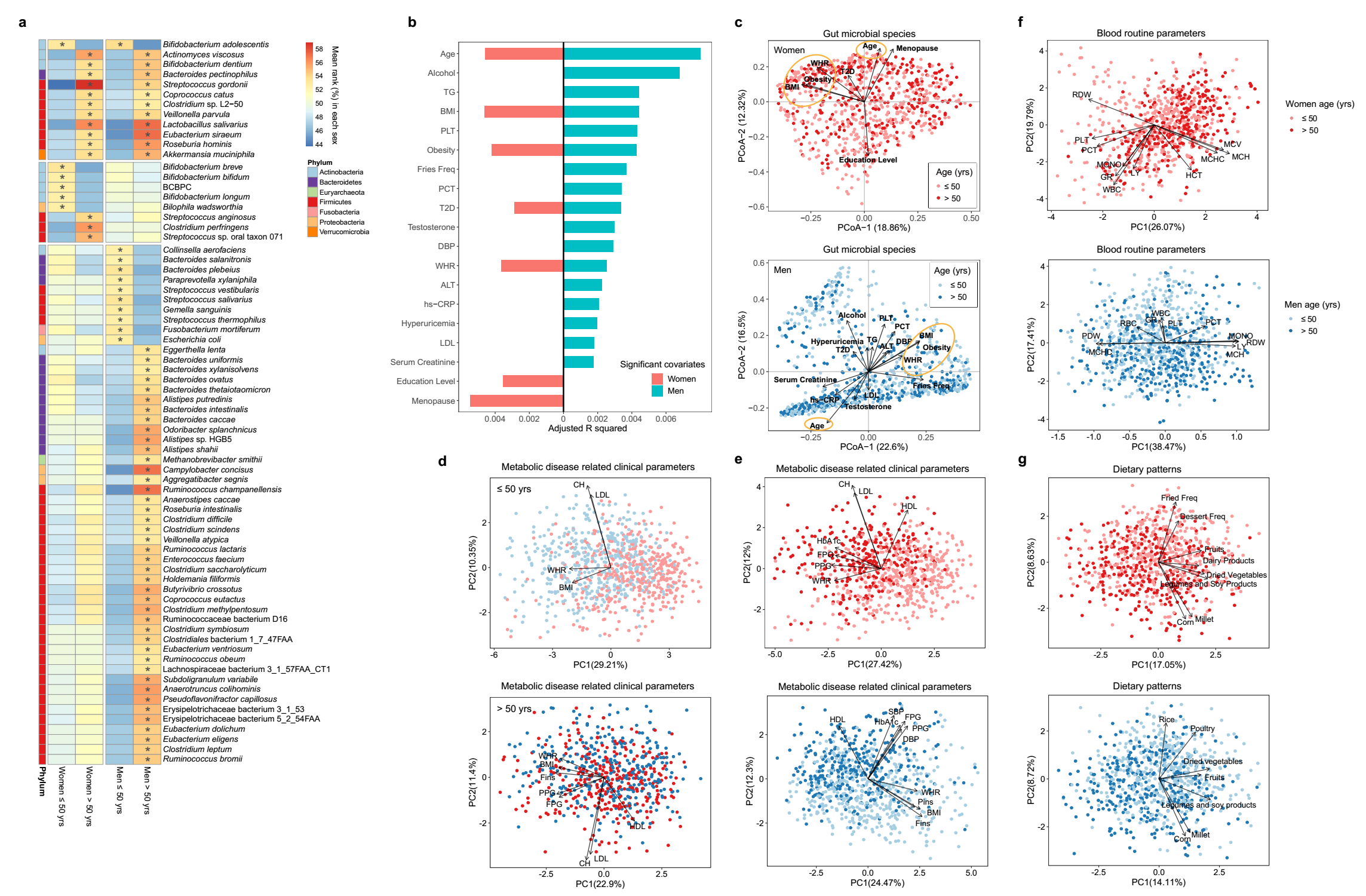

**Extended Data Fig 7. Age- and sex- related differences in the gut microbiota, host phenotypes and their associations in the PG cohort.**

**a,** Heatmap showing the differentially abundant gut microbial common species between the two age groups in women (n=20) and in men (n=64). Wilcoxon rank test, * adjusted *P* < 0.05.

**b,** Horizontal bars showing the amount of inferred variance (adjusted R-squared) explained by each identified significant covariate as determined by PERMANOVA with Bray-Curtis (BC) dissimilarities based on differentially abundant gut microbial species between the two age groups in women (red) and in men (blue). PERMANOVA, adjusted *P* < 0.05.

**c,** Unconstrained PCoA using the BC dissimilarities described in panel (b). Arrows indicate the dimensions of identified significant covariates.

**d,** PCA of blood routine parameters (left), of metabolic parameters (middle) and of dietary patterns (right) measured/collected in women (up) and men (down). For each panel, factors with absolute component scores for PC1 or PC2 ≥ 0.3 are shown as major contributors. Colors of each dot in (c) and (d) indicate adults under (light color) and over 50 years (dark color).

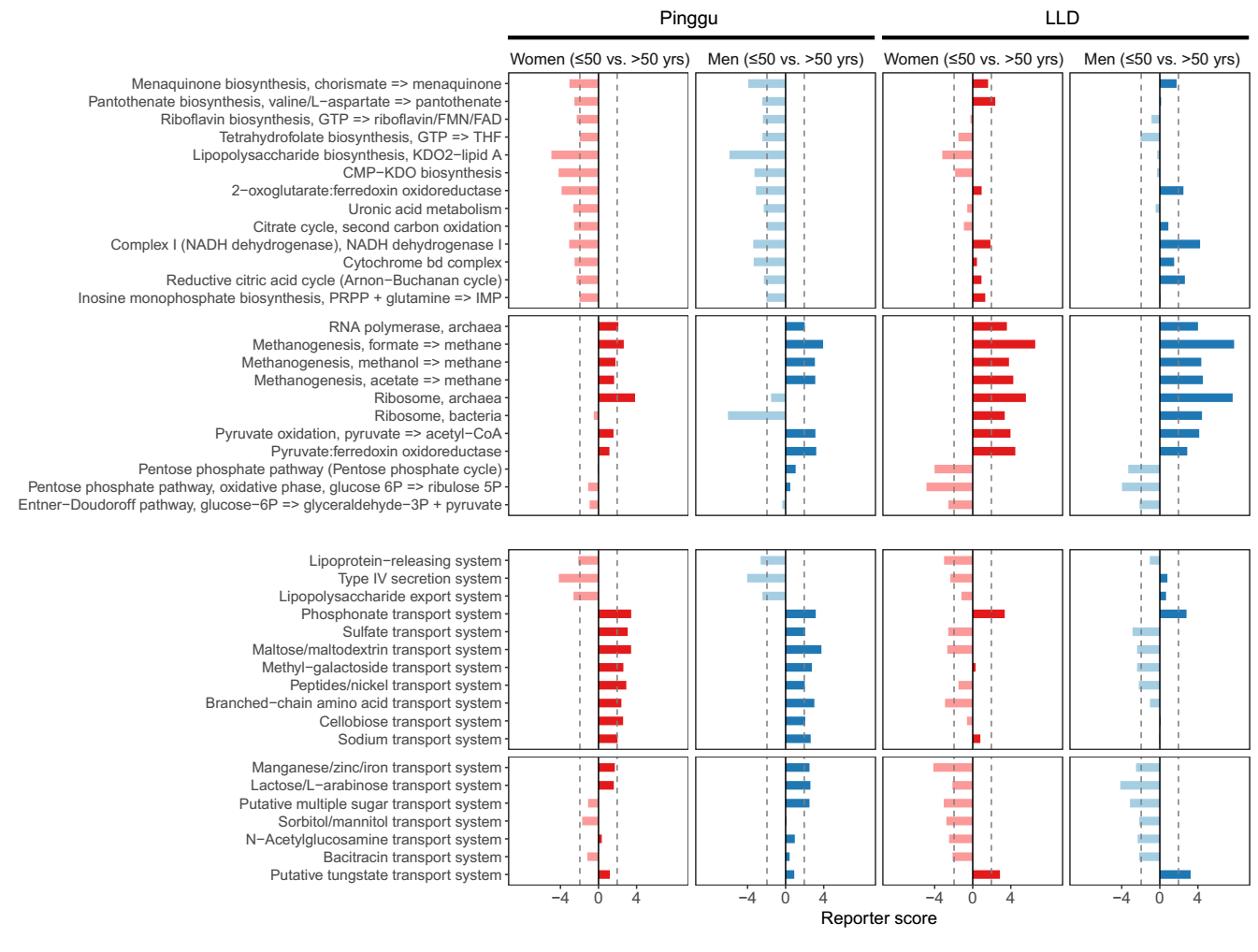

**Extended Data Fig. 8 Gut microbial functional differences between adults below and above 50 years in each sex.**

Differentially abundant modules in adults below (<=50 years, light color) and above 50 years (>50 years, dark color) for women (red) and men (blue) in the PG (left) and LLD (right) cohort, with dashed line indicating a reporter score of 1.96 which corresponding to 95% confidence in a normal distribution. See **Supplementary Table 16-17** for full information.

**Supplementary Tables**

**Supplementary Table 1:** Description and summary statistics of the Pinggu cohort (n=2,338)

**Supplementary Table 2:** Impacts of drugs on the gut microbiota

**Supplementary Table 3:** Description and summary statistics of the Pinggu analysis cohort (n=1,741)

**Supplementary Table 4:** Association of phenotypes with Bray-Curtis dissimilarities at the gene level

**Supplementary Table 5:** Association of phenotypes with Bray-Curtis dissimilarities at the KO level

**Supplementary Table 6:** Association of phenotypes with Bray-Curtis dissimilarities at the common species level

**Supplementary Table 7:** Comparison of gut microbial compositional features between the two sexes

**Supplementary Table 8:** Comparison of the composition of serum bile acids between the two sexes

**Supplementary Table 9:** Associations between phenotypes and microbial features in women (Spearman's correlation)

**Supplementary Table 10:** Associations between phenotypes and microbial features in women (Partial Spearman's correlation adjusting for age)

**Supplementary Table 11:** Associations between phenotypes and microbial features in women (Partial Spearman's correlation adjusting for age and BMI)

**Supplementary Table 12:** Associations between phenotypes and microbial features in men (Spearman's correlation)

**Supplementary Table 13:** Associations between phenotypes and microbial features in men (Partial Spearman's correlation adjusting for age)

**Supplementary Table 14:** Associations between phenotypes and microbial features in men (Partial Spearman's correlation adjusting for age and BMI)

**Supplementary Table 15:** Comparison of gut microbial compositional features between sexes and age groups

**Supplementary Table 16:** Reporter score of KEGG modules between sexes and age groups in the Pinggu cohort

**Supplementary Table 17:** Reporter score of KEGG modules between sexes and age groups in the LLD cohort

**Supplementary Table 18:** Association of phenotypes with Bray-Curtis dissimilarities in the two age groups at the species level

**Supplementary Table 19:** Details for laboratory measurements

**Supplementary Table 20:** List of published gut metagenomic datasets of Chinese and Dutch adults
